# Wide-area all-optical neurophysiology in acute brain slices

**DOI:** 10.1101/433953

**Authors:** Samouil L. Farhi, Vicente J. Parot, Abhinav Grama, Masahito Yamagata, Ahmed S. Abdelfattah, Yoav Adam, Shan Lou, Jeong Jun Kim, Robert E. Campbell, David D. Cox, Adam E. Cohen

**Affiliations:** Chemical Biology Program, Harvard University, Cambridge, Massachusetts, USA; Biophysics Program, Harvard University, Cambridge, Massachusetts, USA; Division of Health Science and Technology, Massachusetts Institute of Technology, Cambridge, Massachusetts, USA; Department of Molecular and Cellular Biology, Harvard University, Cambridge, Massachusetts, USA; Center for Brain Science, Harvard University, Cambridge, Massachusetts, USA; Department of Chemistry, University of Alberta, Edmonton, Alberta, Canada; J.A. Paulson School of Engineering and Applied Sciences; Department of Chemistry and Chemical Biology Harvard University, Cambridge, Massachusetts, USA; Department of Physics, Harvard University, Cambridge, Massachusetts, USA; Howard Hughes Medical Institute, Harvard University, Cambridge, Massachusetts, USA

## Abstract

Optical tools for simultaneous perturbation and measurement of neural activity open the possibility of mapping neural function over wide areas of brain tissue. However, spectral overlap of actuators and reporters presents a challenge for their simultaneous use, and optical scattering and out-of-focus fluorescence in tissue degrade resolution. To minimize optical crosstalk, we combined an optimized variant (eTsChR) of the most blue-shifted channelrhodopsin reported to-date with a nuclear-localized red-shifted Ca^2+^ indicator, H2B-jRGECO1a. To perform wide-area optically sectioned imaging in tissue, we designed a structured illumination technique that uses Hadamard matrices to encode spatial information. By combining these molecular and optical approaches we made wide-area maps, spanning cortex and striatum, of the effects of antiepileptic drugs on neural excitability and on the effects of AMPA and NMDA receptor blockers on functional connectivity. Together, these tools provide a powerful capability for wide-area mapping of neuronal excitability and functional connectivity in acute brain slices.

## Introduction

All-optical neurophysiology (AON)—simultaneous optical stimulation and optical readout of neural activity—provides a promising approach to mapping neural excitability and synaptic strength across wide regions of brain tissue^1^,2. Recent advances in two-photon (2P) calcium imaging AON *in vivo* have enabled input-output measurements on circuits containing up to ~100 near-surface neurons in small cortical regions.^2–4^ However, most of the intact rodent brain remains inaccessible to optical microscopy, and one would ideally like to perform AON simultaneously on many thousands of neurons across multiple brain regions to map spatial variations in function.

Acute brain slices in principle enable wide-area optical mapping across any brain region. While slicing cuts many long-range connections, the procedure preserves the molecular makeup, electrophysiological properties, and local microcircuitry of the component neurons^5^. This preparation is widely used for studies of molecular influences on neural excitability and synaptic transmission (see, e.g. 6-8). Brain slices typically show little spontaneous activity and obviously lack sensory inputs, so optical mapping in brain slices requires a means to evoke activity. Optogenetic stimulation can directly evoke activity in the measured neurons, or can activate axon terminals—even when the axons have been severed from the cell bodies—and evoke postsynaptic responses.^9^ Optical readouts of evoked response could reveal the spatial structure of intrinsic neuronal excitability, of synaptic strength, or of local microcircuit dynamics, and molecular influences thereon.

The optical requirements of wide-area AON in brain slice differ from *in vivo*, suggesting that a distinct approach could be warranted. In brain slice there is a benefit to having a very wide field of view to probe many neurons and brain regions simultaneously. Optical sectioning is important to distinguish in-focus cells from background, but depth penetration is not so important because the plane of the slice can expose any brain structure of interest. One may wish to stimulate many thousands of cells simultaneously, a task beyond the capabilities of current 2P stimulation techniques. If one treats cells as units, the spatial resolution must be sufficient to resolve single cells, but need not resolve fine sub-cellular structures. Time resolution must be sufficient to resolve dynamics slice wide, typically < 200 ms for Ca^2+^ imaging. These factors, discussed in detail below, suggest that one-photon (1P) stimulation and imaging may be preferable over the 2P approaches which have been optimized for *in vivo* use. To achieve 1P AON in brain slice one must (a) identify an actuator/reporter pair with good photostability and minimal optical crosstalk under 1P illumination, and (b) implement a 1P optically sectioned wide-area imaging scheme. Here we combine molecular and optical engineering to address these challenges.

Red-shifted channelrhodopsins have been combined with a GCaMP Ca^2+^ indicator for 2P AON *in vivo*,^10–12^ but 1P GCaMP excitation causes spurious channelrhodopsin excitation. Lower optical crosstalk is achieved by pairing a blue-shifted channelrhodopsin with a red-shifted reporter^13^. Red genetically encoded Ca^2+^ indicators (RGECIs) now offer good sensitivity, but their combination with optogenetic stimulation has been hampered by blue-light induced photoswitching of the mApple-based chromophores used in the most sensitive RGECIs^14–16^. Furthermore, blue channelrhodopsins such as ChR2(H134R) retained some excitation at the yellow (561 nm) wavelengths used to excite RGECIs, introducing crosstalk of the imaging light into the stimulation channel. A truly orthogonal 1P actuator/RGECI reporter pair has not previously been reported.

TsChR, derived from *Tetraselmis striata*,^17^ is the most blue-shifted channelrhodopsin reported to-date, but its initial characterization yielded a poor photocurrent. To our knowledge, TsChR has not previously been used in any optogenetic experiments. Here we show that a version with improved trafficking, eTsChR, drives robust spiking in cultured neurons and in tissue. Combination of eTsChR with a nuclear-localized red-shifted Ca^2+^ reporter, H2B-jRGECO1a, achieved 1-photon AON in cultured neurons and in slice. The blue light used to activate the eTsChR was dim enough to avoid jRGECO1a photoswitching, and the yellow light used to excite jRGECO1a did not spuriously activate the eTsChR.

On the imaging front, 1P structured illumination microscopy (SIM) techniques can achieve optical sectioning in tissue ^18^. We developed a generalized SIM technique based on Hadamard-Walsh encoding and implemented it in a mesoscope imaging system. Hadamard microscopy provides better rejection of out-of-plane fluorescence than do other SIM techniques and offers the ability to make systematic tradeoffs between background rejection and time resolution.

By applying 1P optogenetic stimulation and Hadamard microscopy to acute slices expressing eTsChR and H2B-jRGECO1a, we obtained simultaneous functional characterization of > 6,000 neurons, spread over a region 2.3 × 2.3 mm with 5.6 Hz time resolution. Maps of optically induced activity highlighted distinct cortical layers, which otherwise appeared homogeneous in their fluorescence. We used the AON system to map with cellular resolution the effects of anti-epileptic drugs on neural excitability, and to study cortico-cortico and cortico-striatal functional connectivity. The combined molecular and optical tools provide a powerful system for wide-area investigations of neural function in brain tissue.

## Results

### A spectrally orthogonal Ca^2+^ sensor and channelrhodopsin for 1-photon AON

AON requires a spectrally orthogonal optogenetic actuator and activity reporter (**Fig. 1a**). Examination of channelrhodopsin action spectra and Ca^2+^ reporter excitation spectra suggested that the best approach for 1-photon AON was to use a blue-shifted channelrhodopsin and a red-shifted genetically encoded Ca^2+^ indicator (RGECI). We thus set out to identify protein pairs suitable for this purpose.

**Figure 1:**
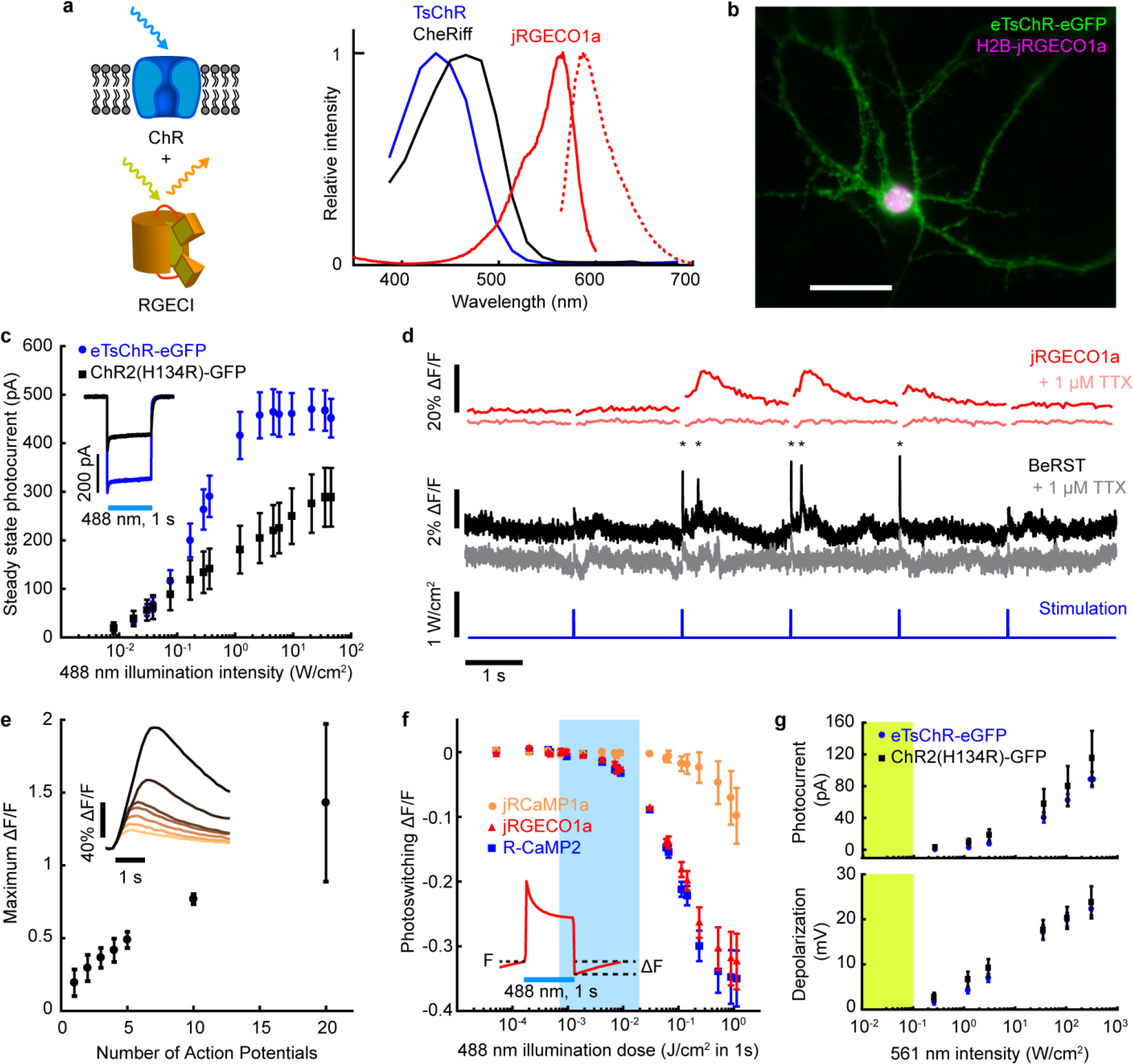
All-optical neurophysiology with a blue-shifted channelrhodopsin and a red-shifted Ca^2+^ indicator. (**a**) Left: Schematic of a spectrally orthogonal channelrhodopsin and RGECI. Right: action spectra of proteins used in this work. Spectra are reproduced with permission from Dana, *et al. eLIFE* (2016) and Klapoetke, *et al. Nat. Meth.* (2014). (**b**) Cultured hippocampal neuron coexpressing H2B-jRGECO1a (magenta) and eTsChR (green). Scale bar 10 μm. (**c**) Steady state photocurrents of eTsChR and ChR2(H134R) in cultured neurons held at −65 mV (1 s pulses, 488 nm, n = 6 cells for each construct). Inset: photocurrent response to 2 W/cm^2^ 488 nm illumination. (**d**) Optogenetic stimulation induced action potentials and corresponding fluorescence transients in a cultured neuron expressing jRGECO1a and eTsChR. Pulses of blue light (488 nm, 10 ms, 680 mW/cm^2^) drove action potentials (*), which were identified via fluorescence of a far-red voltage-sensitive dye, BeRST1 (1 μM, black)^19^. Fluorescence transients of jRGECO1a accompanied action potentials (red). TTX (1 μM) silenced activity in both the voltage (pink) and Ca^2+^ (grey) channels, confirming that signals arose from neural activity and not optical crosstalk. Voltage imaging was performed at 500 Hz with 0.7 W/cm^2^ 640 nm light and calcium imaging was performed at 20 Hz with 1.1 W/cm^2^ 561 nm light. (**e**) Maximum ΔF/F of H2B-jRGECO1a fluorescence vs. number of evoked action potentials in cultured neurons, stimulated via current injection (*n* = 3 cells). Inset: example responses to increasing numbers of action potentials. (**f**) Blue-light induced photoswitching of RGECIs in HEK293T cells under basal Ca^2+^ levels. RGECI fluorescence was recorded at 50 Hz with illumination at 561 nm, 80 mW/cm^2^. Blue illumination (1 s pulses, 488 nm) was added to the yellow illumination. Photoswitching was quantified as the decrease in fluorescence following blue light illumination compared to the initial fluorescence (inset), *n* = 3 FOV, ~50 cells/FOV, for each construct. (**g**) Activation of channelrhodopsins as a function of 561 nm illumination intensity. Top: Steady state photocurrents in cultured neurons voltage clamped at −65 mV. Bottom: Voltage depolarization under current-clamp with an initial potential of −65 mV. Yellow bar indicates typical jRGECO1a imaging intensities. Acquired from cultured rat hippocampal neurons, *n* = 6 for each construct. All error bars indicate mean +/− s.e.m..

We began by comparing the single action potential responses of RGECIs in cultured neurons. jRGECO1a was the most sensitive (ΔF/F = 54 ± 10%, *n* = ~120 neurons), followed by R-CaMP2 and jRCaMP1a, consistent with previous reports (**Supplementary Fig. 1a-b, Supplementary Table 1**)^15^. R-CaMP2 had the fastest kinetics (τ_on_ = 26 ± 10 ms, τ_off_ = 270 ± 20 ms, *n* = ~120 neurons), followed by jRGECO1a (τ_on_ = 47 ± 1 ms, τ_off_ = 440 ± 40 ms, *n* = ~120 neurons) and jRCAMP1a (**Supplementary Fig. 1a-b**, **Supplementary Table 1**). In HEK293T cells, under basal Ca^2+^ conditions, jRGECO1a had the longest photobleaching time constant (τ_bleach_ = 81 ± 5 s, *I*_561_ = 44 W/cm^2^, *n* = 9 cells), followed by R-CaMP2 and jRCaMP1a (**Supplementary Table 1**). Under typical imaging conditions (*I*_561_ = 0.1 W/cm^2^), photobleaching of jRGECO1a was thus < 10% during 1 hr of continuous imaging. While photobleaching is often a concern for 1P imaging, these results established that this effect was minor for wide-area imaging of jRGECO1a. We selected jRGECO1a for its superior sensitivity and photostability.

mApple-based fluorescent sensors, including jRGECO1a, are known to undergo photoswitching under blue light illumination^14^,16. We thus sought a blue-shifted channelrhodopsin that could drive spikes in jRGECO1a-expressing neurons at blue intensities low enough to avoid optical crosstalk. TsChR is the most blue-shifted published ChR (**Fig. 1a**), but was initially reported to produce only ~40% as much photocurrent as ChR2(H134R)^17^ and so has not previously been used in optogenetic applications. Addition of a K_ir_2.1 trafficking sequence (TS) and a GFP expression tag to TsChR led to excellent trafficking in cultured neurons (**Fig. 1b**). We called this construct eTsChR-eGFP. Compared to ChR2(H134R), eTsChR had higher steady state photocurrents (470 ± 42 vs. 288 ± 60 pA, *p* = 0.034, Student’s *t*-test, *n* = 6 neurons each), higher maximum steady state photocurrent densities (13.2 ± 1.2 pA/pF vs. 7.8 ± 2.0 pA/pF, *p* = 0.044, Student’s *t*-test, *n* = 6) and faster on- and off- kinetics (**Fig. 1c, Supplementary Fig. 1c-d**). At the highest blue light intensity tested (33 W/cm^2^), ChR2(H134R) passed a steady state photocurrent of 288 ± 60 pA; eTsChR passed the same steady state photocurrent at a 100- fold lower intensity (0.33 W/cm^2^; **Fig. 1c**).

We co-expressed jRGECO1a and eTsChR in cultured rat hippocampal neurons, and used the far-red voltage-sensitive dye BeRST1^19^ as a ground-truth reporter of neural spiking. Flashes of blue light (0.7 W/cm^2^, 10 ms) induced action potentials, reported by BeRST1 fluorescence, and Ca^2+^ transients, reported simultaneously by jRGECO1a fluorescence (**Fig. 1d**). The sodium channel blocker TTX (1 μM) eliminated the light-evoked transients in both the BeRST1 and jRGECO1a fluorescence channels, confirming that the jRGECO1a response reflected spiking-dependent Ca^2+^ influx and that the optogenetic stimulation did not induce detectable photo-artifacts in the jRGECO1a fluorescence.

Cytoplasmic expression of jRGECO1a in brain slices led to a high level of fluorescence background from reporter present in neuropil, even with the optically sectioned imaging approaches described below. To facilitate imaging in tissue, we fused jRGECO1a to a Histone-2B (H2B) tag to localize expression to the nucleus (**Fig. 1b and Supplementary Fig. 1f**), as previously done for zebrafish^20^ and rat^21^ brain imaging. The nuclear-localized H2B-jRGECO1a showed clearly resolved nuclei with little background between the cells (**Supplementary Fig. 1e**). In cultured neurons, H2B-jRGECO1a responded to single action potentials with good sensitivity (ΔF/F = 19.4 ± 5.3%, *n* = 3 cells), but with slower kinetics than the cytosolic reporter, (τ_on_ = 167 ± 27 ms, τ_off_ = 1,400 ± 270 ms) consistent with previous measurements of nuclear Ca^2+^ dynamics (**Fig. 1e**)^22,23^.

We tested for optical crosstalk between actuator and reporter channels in cells co-expressing the optimized AON constructs. Due to the high sensitivity of eTsChR, the blue light doses needed to elicit spikes (0.7 W/cm^2^ for 10 ms, λ = 488 nm) induced minimal photoartifact in either cytoplasmic or nuclear jRGECO1a compared to a single-spike Ca^2+^ signal (−2% photoartifact in **Fig. 1f** vs. 19% spike response in **Fig. 1e**, **Supplementary Fig. 1h**). Crosstalk from direct blue light excitation of jRGECO1a fluorescence was avoided in the experiments below by interleaved optogenetic stimulation and fluorescence imaging.

The yellow light used for Ca^2+^ imaging (λ = 561 nm, 0.1 W/cm^2^) induced in eTsChR a steady-state photocurrent less than 0.5 pA (**Fig. 1g**), far too small to trigger spurious action potentials. Expression of eTsChR did not significantly affect neurons’ membrane resistance, membrane capacitance, or resting potential compared to controls (**Supplementary Table 2**). Together, eTsChR and H2B-jRGECO1a formed a suitable actuator/reporter pair for crosstalk-free 1P AON.

### Hadamard microscopy enables optical sectioning in ultra-widefield images of acute brain slices

We next sought to perform wide-area optically sectioned imaging of the AON constructs in acute brain slices. To achieve high light collection efficiency over a wide field of view (FOV), we designed a microscope system around a low magnification high numerical aperture objective (Olympus MVPLAPO 2 XC, NA 0.5). In wide-field epifluorescence mode, this microscope imaged a 4.6 mm FOV, large enough to capture most of a hemisphere of a coronal brain slice, with nominal 2.25 μm lateral resolution set by the pixel size on the sCMOS detector. Apart from the optical filters and the mechanical mounts, all elements of the microscope were off-the-shelf components (**Methods**).

To achieve optical sectioning over a wide FOV, we developed a structured illumination approach based on Hadamard encoding. We placed a digital micromirror device (DMD) in the illumination path to enable arbitrary spatiotemporal patterning of the fluorescence excitation. Each DMD pixel mapped to 6.3 μm in the sample plane. The DMD modulated the excitation light with a series of binary illumination patterns such that neighboring sample locations were illuminated with orthogonal intensity sequences (P_1_, P_2_, …, P_n_ in **Fig. 2a**). Raw data consisted of a series of images (F(t_1_), F(t_2_), …, F(t_m_) in **Fig. 2b-1**) acquired with each illumination pattern, which were then demodulated to yield images of the scattered light for each illumination location (**Fig. 2b-2**). Software binary masks then rejected scattered light (**Fig. 2b-3**), akin to physical pinholes used in confocal microscopy. The sum of images over all illumination locations yielded an optical section (**Fig. 2b-4**, **Methods**).

**Figure 2.**
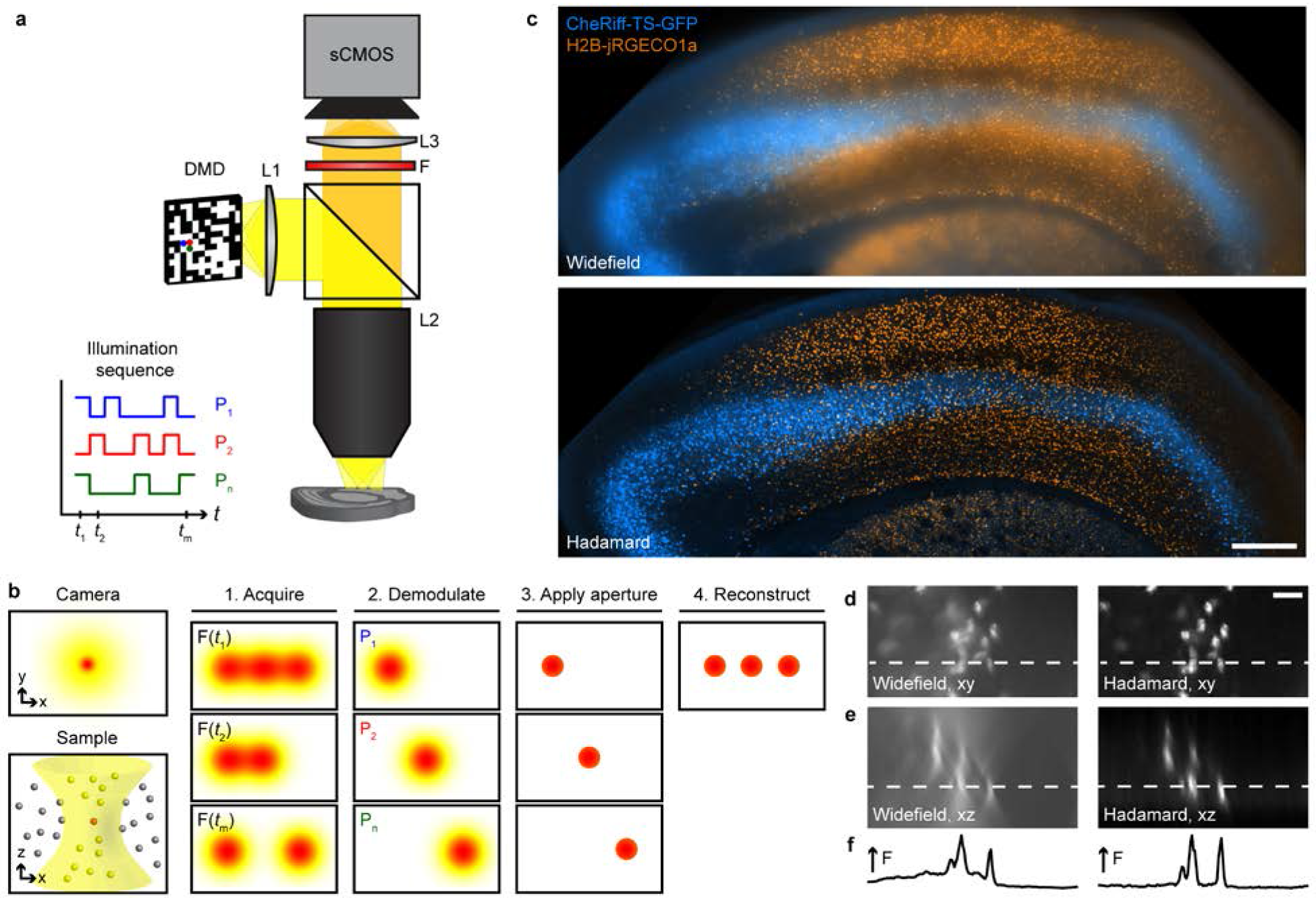
Optical sectioning by Hadamard microscopy. (**a**) Schematic of ultra-widefield microscope, showing orthogonal illumination sequences in neighboring DMD pixels (P_1_, P_2_, …, P_n_). (**b**) Left: In a thick, scattering sample, the in-focus light (red) is dispersed by scattering and mixed with out-of-focus light (yellow). Right: Hadamard microscopy protocol. (1) The sample is illuminated with orthogonal functions at adjacent points. (2) The images are demodulated by matched filtering with the illumination sequence. (3) Scattered light is rejected by a software aperture. (4) The optically sectioned image is reconstructed from a sum of the demodulated images. (**c**) Two-color fluorescence maximum-intensity projection acquired from acute brain slice expressing H2B-jRGECO1a and membrane targeted CheRiff-TS-GFP expressed in L5 pyramidal cells. Top: wide-field epifluorescence. Bottom: Hadamard image. Scale bar 500 μm. (**d-e**) Comparison of widefield and Hadamard image planes in a fixed brain slice expressing membrane-targeted mCitrine illustrating Hadamard background rejection and an improved point-spread function. Scale bars 50 μm. (**f**) Intensity profile from dotted line in (d-e).

To make all projected DMD pixels mutually orthogonal would require prohibitively long digital codes (~10^6^ samples), but because light scatter is mostly local, repeating the codes periodically at separations larger than the scattering point-spread function resulted in minimal crosstalk (**Supplementary Fig. 2a**). Residual crosstalk between repeated codes was scrambled by inverting the sequence of a randomly selected 50% subset of the pixels (**Supplementary Fig. 2a**, Methods). This procedure resulted in series of patterns with 50% duty cycle, uniform mean illumination across the sample, and uniform spatial and temporal spectral density. By varying the number of frames in the Hadamard sequence, one can systematically trade time resolution vs. background rejection. The workflow for acquiring and analyzing Hadamard images is summarized in **Supplementary Fig. 2b** and the code is available as **Supplementary Software**.

We compared Hadamard microscopy to two other SIM techniques, stripe SIM^24^ and HiLo^25^, all implemented using the same DMD and optics. Images of 0.2 μm fluorescent beads in agarose were used to estimate the point-spread functions (PSFs) of the three techniques. As, expected, line sections through the three PSFs gave identical lateral (FWHM 2.7 μm) and axial (FWHM 14.0 μm) resolution near the focus (**Supplementary Fig. 2c,d**). For the low-magnification, wide-area implementation described here, the resolution in all three cases was determined by the intersection of the pixel-size-limited DMD illumination and the camera collection PSFs.

The three imaging techniques differed critically in imaging parameters not captured by the FWHM of the PSFs, however. Stripe SIM and HiLo PSFs had out-of-focus conical lobes, a consequence of out-of-focus points emitting along the same rays as in-focus and laterally offset points. These lobes were absent in Hadamard images, where use of multiple illumination patterns resolved ambiguities in assignment of out-of-focus fluorescence. These lobes did not lie along either the lateral or axial line sections through the PSF, so they did not contribute to the PSF dimensions as usually characterized. To capture the effect of these lobes, we integrated the PSF in the transverse (x-y) plane and plotted total signal as a function of axial coordinate (z). For the Hadamard PSF, total fluorescence decayed to 15% of its peak at a defocus of -30 μm, whereas at the same defocus HiLo retained 38% of peak fluorescence and stripe SIM retained 62% of peak fluorescence (**Supplementary Fig. 2d**).

For the purpose of rejecting out-of-focus background fluorescence in tissue, the integrated transverse fluorescence, not the more commonly used axial line section, is the critical parameter. Thus we expected that Hadamard microscopy would perform better than stripe SIM or HiLo in resolving single-cell signals in densely expressing tissues.

We compared the performance of the three structured illumination techniques in brain tissue (**Supplementary Fig. 3**). The sample comprised an acute 300 μm-thick coronal brain slice, expressing nuclear-targeted jRGECO1a throughout cortex and striatum, and membrane-targeted CheRiff-GFP restricted by an Rbp4-Cre driver to a subset of Layer 5 (L5) pyramidal cells. Hadamard images clearly resolved individual cells, whereas wide-field epifluorescence did not (**Fig. 2c**). In the stripe SIM and HiLo images, out of focus nuclei appeared as rings, a consequence of the conical lobes on the PSF, which prevented clear separation of single-cell images (**Supplementary Fig. 4a**). Light scattering caused the Hadamard signal to decay as a function of image depth with a length constant of 27 μm in acute brain slices (**Supplementary Fig. 4d**) and 113 μm in fixed slices. The difference in signal attenuation was attributed to decreased light scattering after the fixation process.

To quantify the ability of Hadamard microscopy to resolve single-cell signals, we used high-resolution confocal microscopy to make ground-truth maps of the spatial distribution of nuclei in fixed slices densely expressing nuclear jRGECO1a. We then simulated Hadamard images of these cells, using the ground-truth nuclei locations and dimensions, the measured point spread function (**Supplementary Fig. 2c**), and the measured signal decay with depth in acute slice. This procedure yielded estimates of the spurious contribution from all other cells to the fluorescence signal ascribed to each nucleus. In cortical layer 2/3, only 10% of the cells received more than 20% crosstalk from other cells. The crosstalk was lower in other brain regions (**Supplementary Fig. 4c**). Cell nuclei had a stereotyped round and localized shape. The degree of crosstalk could be estimated on a cell-by-cell basis via shape deviations. If desired, cells with crosstalk beyond a threshold value could be discarded from the analysis, though this procedure was not used here. Hadamard microscopy thus enabled optically sectioned imaging with single-cell resolution over wide fields of view in acute brain slices.

**Supplementary Fig. 2e** and the **Supplementary Discussion** compare the shot noise and technical noise properties of Hadamard and other SIM techniques. Hadamard performed as well as or better than the other techniques by these parameters.

### Mapping excitability in acute slices

To map neural excitability, we applied Hadamard microscopy with simultaneous optogenetic stimulation in acute mouse brain slices expressing the actuator-reporter pair. We co-injected AAV9-hSyn-DO-H2B-jRGECO1a and AAV9-hSyn-eTsChR in cortex and striatum of wild-type P0-2 mouse neonates (**Fig. 3a**). Both proteins expressed well and were readily visualized via Hadamard imaging in 300 μm acute brain slices from 3-week-old animals (**Fig. 3b-c**). We performed Hadamard AON measurements in a region 2.3 × 2.3 mm, set by the size of the expressing region. Cell signals were acquired from a depth of 32.2 ± 12.7 μm (**Supplementary Fig. 4b**).

**Figure 3:**
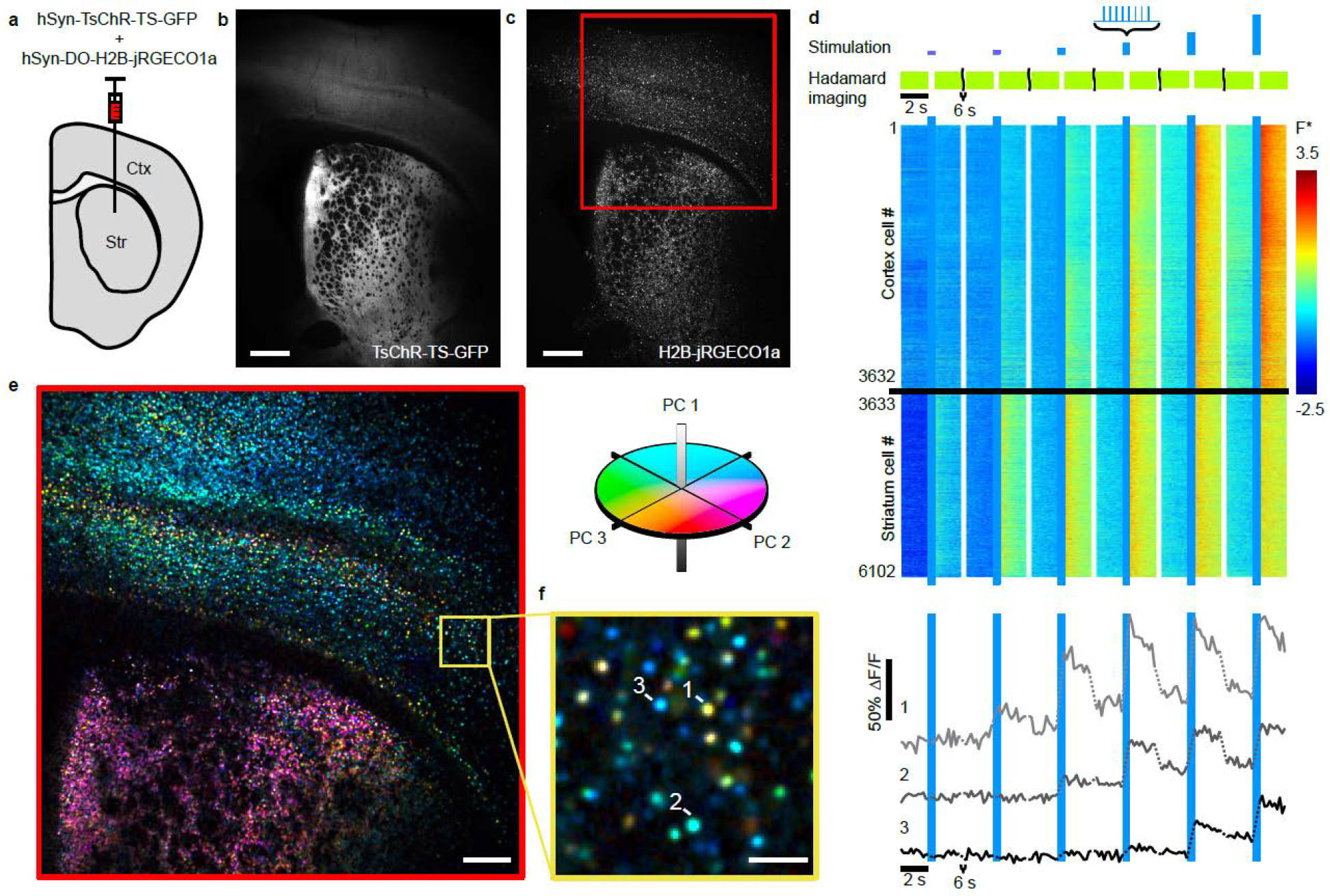
Ultra-widefield AON in acute brain slices. (**a**) AAV9 viruses coding for hSyn-eTsChR and hSyn-DO-H2B-jRGECO1a were co-injected in neonatal mouse cortex and striatum. (**b**) Maximum intensity projection of a Hadamard z-stack of eTsChR expression in a coronal corticostriatal slice from a P21 mouse. (**c**) Same as (b) in the H2B-jRGECO1a channel. Functional data were acquired within the red square. (**d**) Top: Stimulation and imaging protocol. The highlighted FOV in (c) was stimulated with eight 5 ms pulses of 488 nm light at 20 Hz with intensities of 15, 30, 60, 120, 240, and 480 mW/cm^2^. Bottom: heat map of 6,102 single-cell fluorescence traces extracted from the slice in (b-c). Individual fluorescence intensities traces were normalized as F* = (F - mean(F))/std(F). White breaks separate measurements at different optogenetic stimulus intensities. (**e**) Left: boxed region in (c) with cells colored by the principle component amplitudes of the functional responses. See **Methods** for additional details. (**f**) Left: close-up of the yellow boxed region of (e). Right: three example single-cell fluorescence traces. Dotted lines indicate pauses in Hadamard imaging (400 ms during optogenetic stimulation, 6 s between stimuli). ΔF/F is defined relative to the intensity in the first imaging epoch. Scale bars 250 μm in (b, c, e) and 50 μm in (f).

To probe excitability, we exposed the slice to a series of wide-field blue stimuli of increasing strength, interleaved with Hadamard imaging of H2B-jRGECO1a with yellow light (561 nm, 100 mW/cm^2^, **Fig. 3d**). Hadamard images were first acquired for 2 s to establish baseline fluorescence. Then a brief burst of blue light pulses (470 nm, 8 pulses, 15 mW/cm^2^, 5 ms duration, 20 Hz) evoked neural activity, followed by another 2 s of Hadamard imaging to record the response. This image-stimulate-image procedure was repeated at 10 s intervals, six times, with the intensity of the blue light doubling upon each repetition to a maximum of 480 mW/cm^2^. This measurement protocol reported the changes in intracellular Ca^2+^ concentration as a function of optogenetic stimulus strength.

We used a 2D peak-finding algorithm to identify *n* = 6,102 responding cells in the Hadamard images of a single brain slice (**Fig. 3d**). Neighboring cells often showed distinct patterns of Ca^2+^ dynamics, while interstitial regions showed undetectable fluorescence (**Supplementary Fig. 5a,b**), confirming that Hadamard microscopy effectively rejected scatter and out-of-focus background. The yellow light used for Ca^2+^ imaging induced spurious activity in only 0.46 ± 0.03% of cells (*n* = 38,835 cells, 9 slices, **Supplementary Fig. 5c,d**), establishing that the imaging light only weakly activated eTsChR. The sodium channel blocker tetrodotoxin (TTX, 1 μM) abolished blue light evoked responses slice wide, confirming that Ca^2+^ responses were due to action potential firing (**Supplementary Fig. 5e,f**) and, furthermore, that blue light-induced photoswitching was minimal.

We tested the long-term stability of the preparation. The optogenetically induced Ca^2+^ signal was stable over a 78 minute session comprising 7 repeated imaging cycles (**Supplementary Fig. 5g,h**). During this period the population-average optically evoked ΔF/F at the strongest stimulus decreased modestly from 64 ± 0.7% to 52 ± 0.7%, *n* = 3,195 cells. These results demonstrate the capability for repeated measurements over > 1 h in a single sample.

Cells showed different patterns of response in the striatum vs. cortex, but we also observed cell-to-cell variability within the cortex. To characterize this variability, we applied principal components analysis (PCA) to a set of single-cell recordings. First, we repeated the excitability measurement on 9 slices from 2 animals, recording from a total of *n* = 32,103 cells across cortex and striatum. Measurement runs (comprising six measure-stimulate-measure sequences) were repeated at 5 minute intervals, 3 times per slice. PCA identified 3 main temporal components in the single-cell fluorescence responses (**Supplementary Fig. 5i**, **Supplementary Methods**). Examination of the PC temporal waveforms showed that PC1 measured overall fluorescence response amplitude, PC2 captured a left-right shift in the sigmoidal excitability profile, and PC3 largely captured a stimulus-dependent increase in baseline fluorescence.

We then decomposed the fluorescence waveform at each pixel into its principal components (PCs), and color-coded each pixel by its PC amplitudes (**Fig. 3e**, **Supplementary Fig. 5j**, **Supplementary methods**). Despite coloring each pixel independently, individual cells appeared homogeneously colored in the resulting image (**Fig. 3f**), consistent with the low cell-to-cell fluorescence crosstalk. These maps revealed striking colored bands running along the cortical layers, demonstrating different functional responses in different brain regions. Intriguingly, some layers appeared relatively homogeneous (L2/3, L4, L6), while cells in L5 had larger cell-to-cell variations in response. These results demonstrate that Hadamard AON can map excitability over large areas of acute brain slice with single cell precision.

### Mapping pharmacological responses with Hadamard AON

Wide-area AON offers a means to map the cell type and region-specific effects of pharmacological or other perturbations on neural excitability. We performed excitability measurements on acute slices before and after applying the antiepileptic drugs (AEDs) retigabine (25 μM), carbamazepine (100 μM), and phenytoin (100 μM). To quantify the drug effect, we measured the pixel-by-pixel change in mean amplitude, ΔF, of the optogenetically induced response—a parameter close to the first principal component that emerged from the unsupervised analysis above. Each drug had different effects in striatum and cortex, and attenuated cortical excitability in a distinctive spatial pattern (**Fig. 4a**).

**Figure 4:**
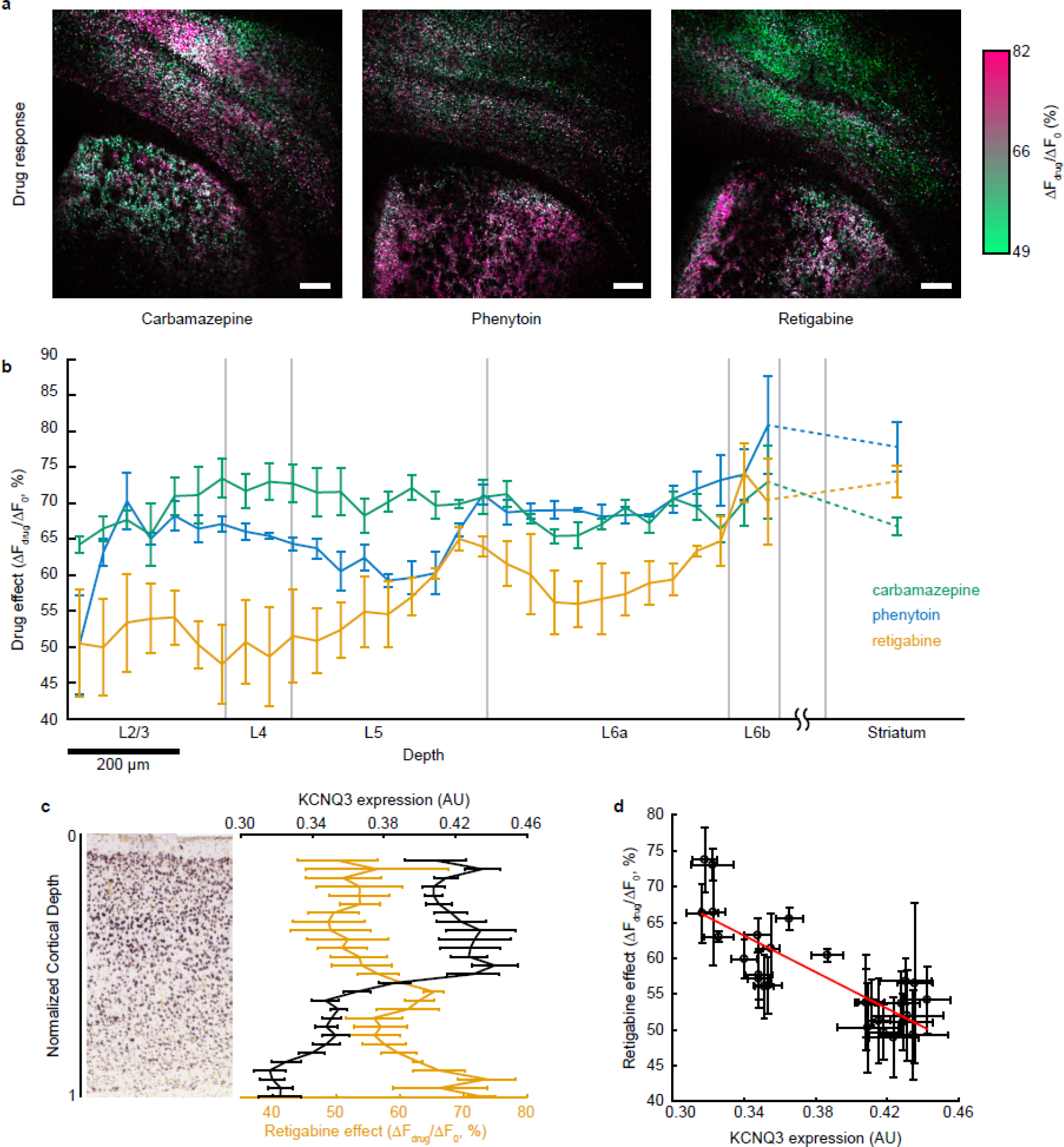
Mapping effects of anti-epileptic drugs (AEDs) on excitability. (**a**) Maps of AED effects on excitability. Slices were measured using the excitability protocol as in Fig. 3. The protocol was repeated five times before drug addition and four times after addition of carbamazepine (100 μM), phenytoin (100 μM), or retigabine (25 μM). The ratio of mean optogenetically induced change in fluorescence for each cell before (ΔF_0_) and after drug addition (ΔF_drug_) is encoded as color in a green to pink axis. Scale bars 250 μm. (**b**) Average drug response (ΔF_drug_/ΔF_0_) as a function of cortical depth for *n* = 3 slices for each drug. All striatal cells in a slice were pooled into a single bin. Data represents *n* = 9,793 cells for carbamazepine, 11,858 cells for phenytoin, and 10,103 cells for retigabine. Error bars represent s.e.m. over *n* = 3 slices for each drug. (**c**) Left: *in situ* hybridization image from Allen Brain Atlas experiment #100041071 showing KCNQ3 expression in somatosensory cortex of a P28 mouse. Right: cortical depth dependence of retigabine drug effect (same as Fig. 4b) and KCNQ3 expression level determined from *in situ* hybridization images of *n* = 11 slices from the Allen Brain Atlas. (**d**) Data from (c) showing effect of retigabine on excitability vs. KCNQ3 expression. Best fit line is indicated in red. Error bars indicate s.e.m., treating each slice as an independent measurement.

We sorted cells into bins based on their cortical depth and visualized mean AED response as a function of cortical depth, averaged over *n* = 3 slices per drug (**Fig. 4b**). Carbamazepine and phenytoin, both sodium channel blockers, showed relatively uniform suppression of excitability as a function of cortical depth, but retigabine showed a graded response, weakest in L6b and strongest in L4.

Retigabine is a specific positive allosteric modulator of K_v_7 channels, and its primary target is thought to be the K_v_7.2/7.3 heteromer^26^, coded for by the genes KCNQ2 and KCNQ3. We examined the Allen Brain Atlas map of the expression level of KCNQ3 ^27^, as determined by RNA *in situ* hybridization (ISH), and found statistically significant correlation between KCNQ3 expression level and effect of retigabine (Pearson’s *r* = -0.40, 95% confidence interval between -0.022 and -0.69 obtained by bootstrapping, **Fig. 4c-d**). Higher expression of KCNQ3 correlated with greater inhibition of excitability by retigabine, as one would expect for a potassium channel activator. An independent ISH study in adult animals reported a similar distribution of KCNQ2 and KCNQ3^28^. These results establish a connection between the Hadamard AON measurements and the underlying pattern of ion channels.

### Probing functional connectivity with ultra-widefield AON

We next sought to extend the Hadamard AON platform to measurements of functional connectivity. Although slicing interrupts many long-range projections, optogenetic stimulation of axon terminals can nonetheless evoke local neurotransmitter release and postsynaptic responses^9^. We reasoned that sufficiently strong presynaptic stimulation would drive postsynaptic spikes, which could be detected via H2B-jRGECO1a.

To achieve this goal, the channelrhodopsin must traffic efficiently to axon terminals. We found that expression of eTsChR was predominantly localized to the soma and dendrites (**Supplementary Fig. 6a**). We thus explored CheRiff-TS-GFP (CheRiff), a blue-light sensitive, high-photocurrent channelrhodopsin^13^. CheRiff trafficked well in axons and was 2.3-fold more sensitive to blue light than eTsChR **Supplementary Fig. 6c-d**). CheRiff was also more sensitive to yellow light, raising the possibility of spurious activation by the 561 nm imaging laser. Under typical imaging conditions (561 nm, 100 mW/cm^2^) CheRiff photocurrent was 0.9% of the maximum photocurrent (95% confidence interval 0.8 to 1%, *n* = 7 cells), whereas eTsChR photocurrent was < 0.1% of its maximum photocurrent (**Supplementary Fig. 6d**).

We designed an experiment to express CheRiff in L5 cortico-striatal neurons following a previously described protocol^29^,30, and to test the postsynaptic response via Ca^2+^ imaging in the striatum. The CheRiff vector comprised CAG-DIO-CheRiff-TS-GFP (Cre-on CheRiff), which we injected into neonatal Rbp4-Cre^+/−^ mice to target expression to a population of excitatory L5 neurons. We concurrently injected hSyn-DO-H2B-jRGECO1a (Cre-off nuclear Ca^2+^ indicator) to drive reporter expression throughout striatum and cortex (**Fig. 5a**).

**Figure 5:**
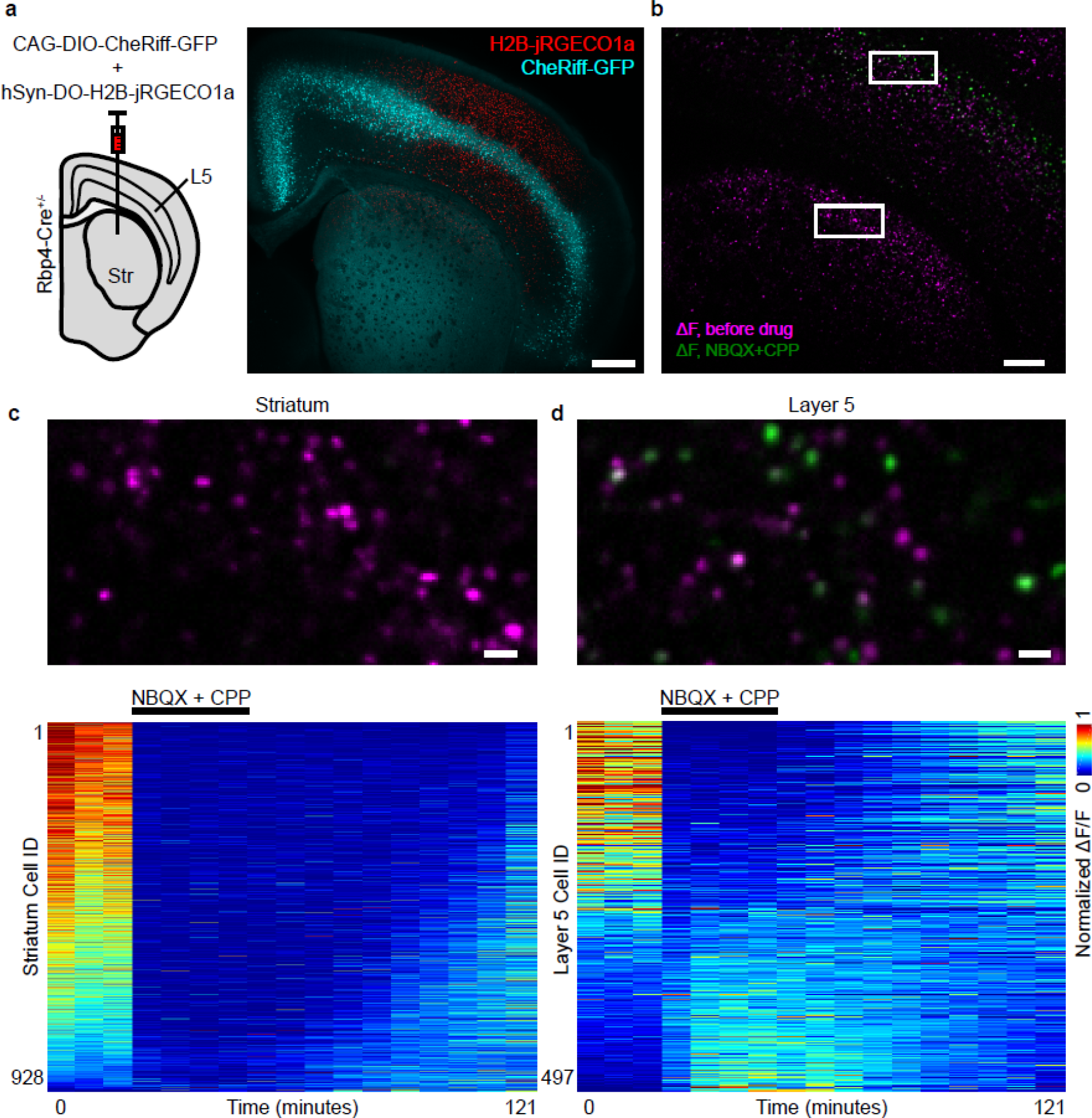
Mapping functional connections. (**a**) Left: viral constructs for mapping functional connections. Cre-dependent AAV9-CAG-DIO-CheRiff-GFP and AAV9-hSyn-DO-H2B-jRGECO1a were co-injected in Rbp4-Cre^+/−^ neonatal mice. Right: at P21, CheRiff-GFP expressed in Cre^+^ L5 pyramidal cells, including corticostriatal projection neurons. H2B-jRGECO1a expressed broadly in cortex and striatum. Images represent a maximum intensity projection of a Hadamard z-stack. Scale bar 500 μm. (**b**) Mean optogenetically induced fluorescence transients, ΔF, before (magenta) and after (green) addition of excitatory blockers, NBQX (10 μM) and CPP (10 μM). Stimulation was performed as in Fig. 3. Images are the median of 3 runs before and 4 runs after adding excitatory blockers. Scale bar 250 μm. (**c**) Magnified views of indicated regions in striatum and Layer 5 in (b). Scale bar 25 μm. (**d**) Mean optogenetically induced fluorescence response, ΔF, for each cell before, in the presence of, and during washout of excitatory blockers. Left: striatum. Right: Layer 5. Each column represents the mean optogenetically induced ΔF of an experimental protocol as in Fig. 3d. The slice was measured over 121 minutes (5-10 minutes between measurements, 22 minutes before last measurement). For visualization, each cell trace was normalized by its mean.

First, we tested the slices for spurious activity elicited by the yellow imaging light. Very few striatal neurons showed a detectable increase in H2B-jRGECO1a signal caused by 561 nm imaging illumination (0.32 ± 0.001%, *n* = 3137 cells, 2 slices, **Supplementary Fig. 6e-h**), confirming that the yellow light did not excite axon terminals enough to drive postsynaptic spikes in most cases. This crosstalk performance is not significantly different from that in the eTsChR-based excitability measurements described above (0.46 ± 0.03%, n = 38,835 cells, 9 slices, *p* = 0.25, two-proportion z-test, **Supplementary Fig. 5c,6h**). In excitability-style measurements with CheRiff, a significantly larger proportion of neurons showed imaging light-induced activation (2.3 ± 0.5%, n = 944 cells, 2 slices, *p* = 8×10^−10^). Thus, the superior axonal trafficking of CheRiff made it the preferred actuator for functional connectivity measurements, while the lower yellow-light crosstalk of eTsChR made it the preferred actuator for excitability measurements.

We then repeated the blue-light stimulation and imaging protocol previously used for excitability measurements while monitoring downstream responses in the striatum. Blue light induced nuclear Ca^2+^ transients across the cortex and striatum (**Fig. 5b**). Blockers of excitatory transmission, NBQX and CPP, reversibly eliminated the responses in the striatum, Layer 6, and Layer 2/3, confirming that these responses were synaptically evoked (**Fig. 5c**) and that there was negligible blue light crosstalk into the fluorescence signals.

To our surprise, addition of NBQX and CPP reversibly increased the optogenetically induced activity in a population of cells in L5 (**Fig. 5d**). The location of these cells amidst the Rbp4 population suggested that these cells expressed both the actuator and reporter (likely a consequence of imperfect silencing of DO-H2B-jRGECO1a in Rbp4-Cre^+^ neurons^31^). The increase in excitability upon excitatory blockade then implies a disinhibitory mechanism, i.e. that these L5 cells received disynaptic inhibition from Rbp4-Cre labeled L5 pyramidal cells, which was relieved under excitatory blockade. The remaining cells in L5 showed a reversible decrease of activity in the presence of excitatory synaptic blockers, similar to the phenotypes in striatum and other cortical layers. These intermixed responses highlight the importance of performing single cell resolution measurements with Hadamard microscopy. Further, although Hadamard microscopy of jRGECO1a can only study supra-threshold responses, these results shown that judicious pharmacological application can dissect a system’s functional connectivity.

## Discussion

Through detailed photophysical characterization of optogenetic actuators and reporters, we identified pairs that can be used in tandem with minimal 1P crosstalk. A pairing of CheRiff and jRCaMP1b was recently demonstrated in cultured neurons, but crosstalk was not measured quantitatively and the genetic constructs were not tested in tissue ^32^. Despite the well reported photophysical blue light artifacts in jRGECO1a, we found that sufficiently sensitive optogenetic actuators could induce neuronal responses at blue light intensities where these artifacts were minimal. The far blue-shifted channelrhodopsin, eTsChR, enabled measurements of intrinsic excitability, and the highly sensitive channelrhodopsin, CheRiff, enabled measurements of functional connectivity, in both cases with minimal crosstalk from the yellow imaging laser. Finally, nuclear localization of the reporter, combined with Hadamard structured illumination microscopy enabled resolution of single-cell signals across wide areas of brain slice. The combined toolbox for 1P AON enables wide-area mapping of excitability and of functional connectivity in brain tissue, and studies on the effects of perturbations thereon.

Questions of where and how neuroactive compounds affect neuronal function are difficult to answer with conventional techniques. Typically, compound distribution is investigated by radiographic labeling experiments. Such results are convolved with possible nonspecific binding of the molecule and with expression of the target in the neuropil, preventing single cell identification. The 1P AON technique provides a high spatial resolution functional alternative to radiographic mapping. We show differential response profiles for three AEDs—one molecularly specific drug, retigabine, whose response profile matched its known target distribution, and two non-specific drugs, carbamazepine and phenytoin. Measurements on other drugs may provide insights into their specific cellular and regional targets. Hadamard AON could also be used to probe the effects of neuropeptides, neuromodulators, hormones, genetic mutations, or environmental perturbations (e.g. temperature, oxygen, metabolites) on brain-wide patterns of neural excitability.

By extending these assays to measurements of functional connectivity, we show that this 1P AON toolbox can be also be used for circuit dissection. The all-optical connectivity assay of Figure 5 shows that Rbp4-Cre positive neurons have a strong excitatory drive across striatum, consistent with previous results ^33^. The net effect of layer 5 stimulation on other cortical layers was not previously well established—most L5 neurons are excitatory but also recruit strong inhibition via parvalbumin and somatostatin neurons across the cortical column ^6^,34. We found a clear net excitatory effect of Rbp4-Cre neuron activation in many cells of L2/3 and L6a of the cortex. Within L5 we found a heterogeneous response, where inhibition outweighed excitation in Rbp4-cre positive neurons (and possibly others which remained nonresponsive during the entire experiment) but excitation outweighed inhibition in other neurons in L5. While this paper was in review, another study interrogated the same circuit with optogenetic stimulation and simultaneous triple whole cell patch clamp, with broadly similar conclusions^35^, though the rigors of patch clamp limited the measurements to a few tens of neurons overall.

Both 1P AON and Hadamard microscopy can be used independently and neither technique is limited to neuroscience applications. The far blue spectrum and excellent sensitivity of eTSChR open the possibility to pair it with red-shifted fluorescent sensors of many other modalities, such as pH, cyclic AMP, or neurotransmitters. The broad spectral range of Hadamard microscopy opens possibilities for high-speed optically sectioned imaging of many different fluorescent reporters, including simultaneous imaging of multiple modalities.

There are many microscopy techniques which could in principle be used for AON in brain slices. Here we briefly outline the factors which led us to develop Hadamard microscopy rather than using an established technique. Spinning disk confocal microscopy^36^ in principle provides high temporal resolution and good optical sectioning, but existing spinning disk optics lack sufficient etendue to capture the FOV and NA of the wide-area objective. One could mimic the function of a spinning disk system by activating individual DMD pixels sequentially in a tiled array, acquiring one image per illumination pattern, and then using software spatial filtering to keep only the in-focus component of each point illumination pattern. This approach would yield the same PSF as Hadamard microscopy. In the **Supplementary Discussion** we show that the Hadamard technique offers better shot-noise limited signal-to-noise ratio for a time- and intensity constrained system. Furthermore, with sparse multi-focal confocal illumination, transient events at un-illuminated pixels would be missed.

As discussed above, stripe SIM and HiLo techniques are alternatives which could be implemented with the same DMD optics as Hadamard microscopy. The improved PSF shape (relative to stripe SIM and HiLo) and the absence of static illumination noise (relative to HiLo) favored Hadamard microscopy. The lower temporal resolution of Hadamard relative to the other SIM techniques did not constrain the ability to map nuclear Ca^2+^ dynamics, though better time resolution may needed for other fluorescent reporters. Improvement in the time resolution of Hadamard microscopy via compressed sensing techniques are anticipated.

2P mesoscopes currently hold the record for most single neurons (~3000) recorded simultaneously in tissue^37^. 2P-mesoscopes have greater depth penetration than SIM techniques, making them more suitable for *in vivo* studies at present. Point-scanning based mesoscopes have achieved pixel rates of ~2×10^7^/s over 0.6 × 0.6 mm FOVs but the requirement to translate the beam long distances limits pixel rates over large FOVs (4.4 × 4.2 mm) to 5.6×10^6^/s. Acousto-optical steering allows fast 2P random-access imaging^38^, but this technique has only been demonstrated in a FOV of 0.5 mm, limited by the etendue of the acousto-optical deflectors. With 12-pattern Hadamard, we achieved comparable data rates of 1.2 ×10^7^/s over a 4.6 × 2.3 mm FOV, with optically sectioned single-cell resolution. With improved control software to synchronize Hadamard patterns to the rolling shutter of the camera, pixel rates of 3.3×10^7^ pixels/s over the entire 4.6 × 4.6 mm FOV would be possible with current camera technology. Finally, in contrast to 2P-mesoscopes, Hadamard microscopy is readily implemented with inexpensive LED or diode laser illumination across a broad range of excitation wavelengths.

For precisely targeted single-cell stimulation, 2P optics are essential, but for wide-area optogenetic stimulation, 1P optics are preferable, as follows: 2P optogenetic stimulation requires time-average optical powers of 20 – 80 mW/cell,^2–4^. Maximal safe steady-state 2P optical power into intact brain tissue is ~200 mW^39^, limiting simultaneous 2P stimulation to at most a few tens of neurons at a time. 1P optogenetic stimulation requires approximately 10^6^-fold lower time-average power (~50 nW/cell)^13^, and thus is readily applied over wide areas of tissue to many thousands of cells simultaneously.

In the present work, the maximum number of neurons recorded in parallel (6,102) was limited not by data rate but by the expression pattern of the virally delivered constructs. In transgenic animals or with recently developed systemic gene delivery techniques^40^, Hadamard microscopy could tile a complete sagittal slice in 7 FOVs, fast enough for functional measurements over acute slices from an entire mouse brain. The resulting brain-wide functional mapping could provide an unbiased approach to studying excitability and functional connectivity.

## Acknowledgments

We thank Vaibhav Joshi, Katherine Williams, and Melinda Lee for technical assistance. We thank Bernardo Sabatini for Rbp4-Cre mice, and Christopher Werley for assistance with the microscope design. We thank Daryl Lim for providing HiLo reconstruction code. Joshua Sanes provided support for the cloning of H2B-jRGECO1a. This work was supported by the Howard Hughes Medical Institute. SLF was supported by an NSF Graduate Research Fellowship. VJP was supported by a Becas Chile scholarship. Work in David Cox’s lab was supported by IARPA (contract #D16PC00002) and the Mind Brain Behavior Faculty Award of Harvard.

## Author contributions

SLF designed, created, and calibrated the optogenetic constructs. VJP designed and built the Hadamard microscope with early assistance from JJK. AG and MY cloned H2B-jRGECO1a and characterized its response via patch clamp electrophysiology. AA cloned eTsChR. YA helped with brain slice work. SL provided brain slices broadly expressing fluorescent proteins. REC supervised the contribution from AA. DDC supervised the contribution from AG. AEC conceived the Hadamard microscopy concept and supervised the research. SLF and VJP acquired data. SLF, VJP, and AEC analyzed data. SLF, VJP, and AEC wrote the paper.

## Data and code availability

Constructs will be made available on Addgene. Code for Hadamard pattern generation and image reconstruction, as well as raw data examples are included as supplementary information.

## Competing financial interests

AEC and VJP have filed a patent application on Hadamard microscopy. AEC is a co-founder of Q-State Biosciences.

**Supplementary Figure 1:**
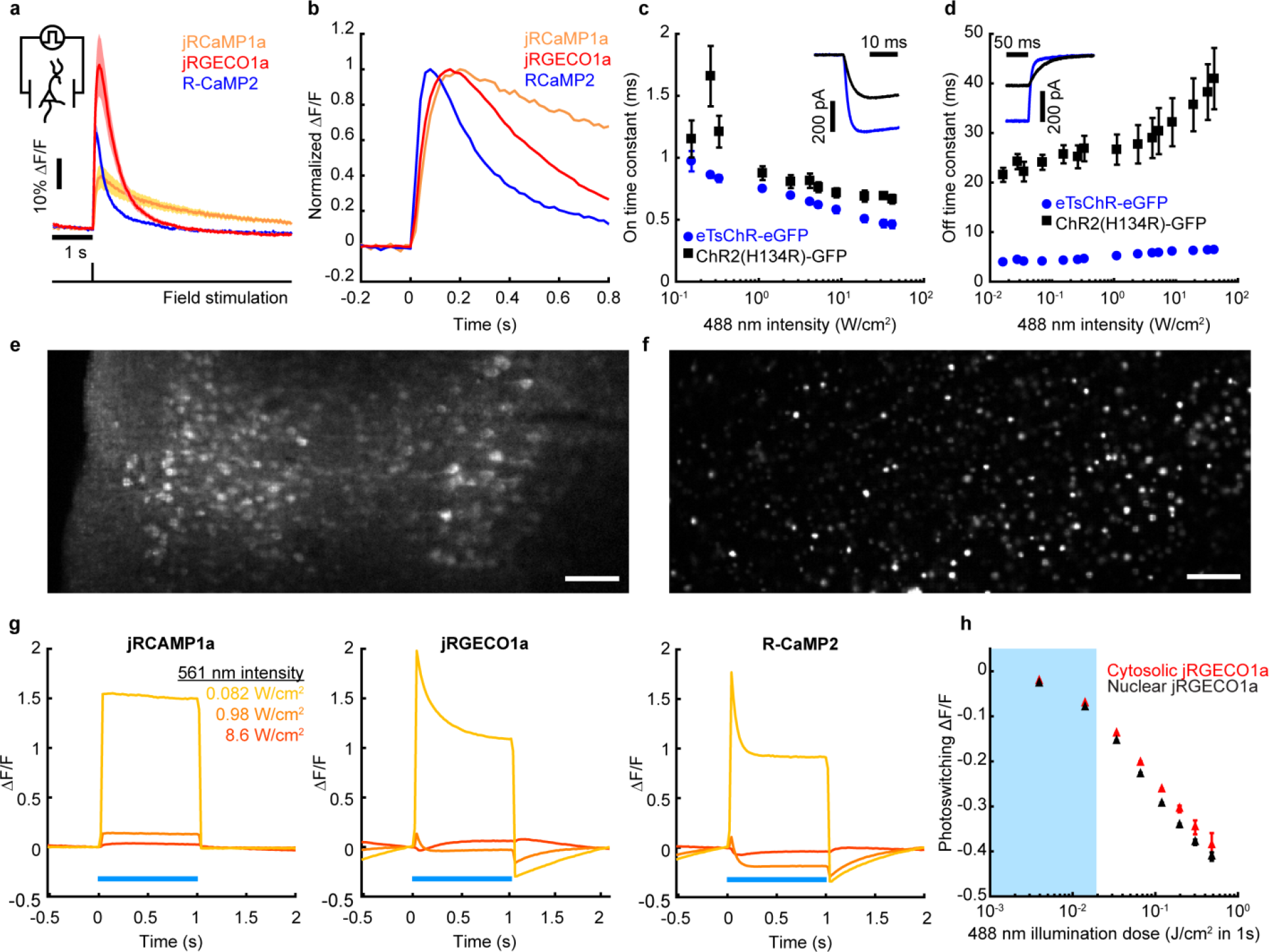
Photophysical characterization of RGECIs and eTsChR. (**a**) Single action potential responses of RGECIs in cultured rat hippocampal neurons. Dark lines indicate the average of 3 FOVs, ~30 cells/FOV, for R-CaMP2 and 4 FOVs for jRGECO1a and jRCaMP1a. Colored bands indicate +/- s.e.m.. Dishes were stimulated with 1 ms field stimulation pulses. RGECI fluorescence was recorded at 50 Hz. (**b**) Kinetics of the RGECIs, shown by plotting data in Fig. 2b normalized to peak ΔF/F. (**c**) Channelrhodopsin activation time constant as a function of 488 nm illumination intensity. Inset: photocurrents during illumination start. (**d**) Closing time constants. Inset: photocurrents during illumination stop. (**e**, **f**) Maximum intensity projections of Hadamard z-stacks from acute cortical slices prepared from mice injected with (e) cytosolic AAV1-syn-NES-jRGECO1a or (f) nuclear-targeted AAV9-syn-DO-H2B-jRGECO1a. Scale bars 100 μm. (**g**) Blue light-induced photoswitching of RGECIs in HEK293T cells. Fluorescence of the RGECI was excited at 561 nm with the indicated illumination intensity. A 1 s pulse of blue illumination (1.1 W/cm^2^ 488 nm light) was added to the yellow illumination (blue bars). In jRGECO1a and R-CaMP2 the blue illumination switched the protein into a state with reduced fluorescence. (**h**) Comparison of photoswitching in cytosolic and nuclear-localized jRGECO1a. Blue bar represents range of illumination doses used for optogenetic stimulation in this study. All error bars indicate s.e.m.

**Supplementary Figure 2:**
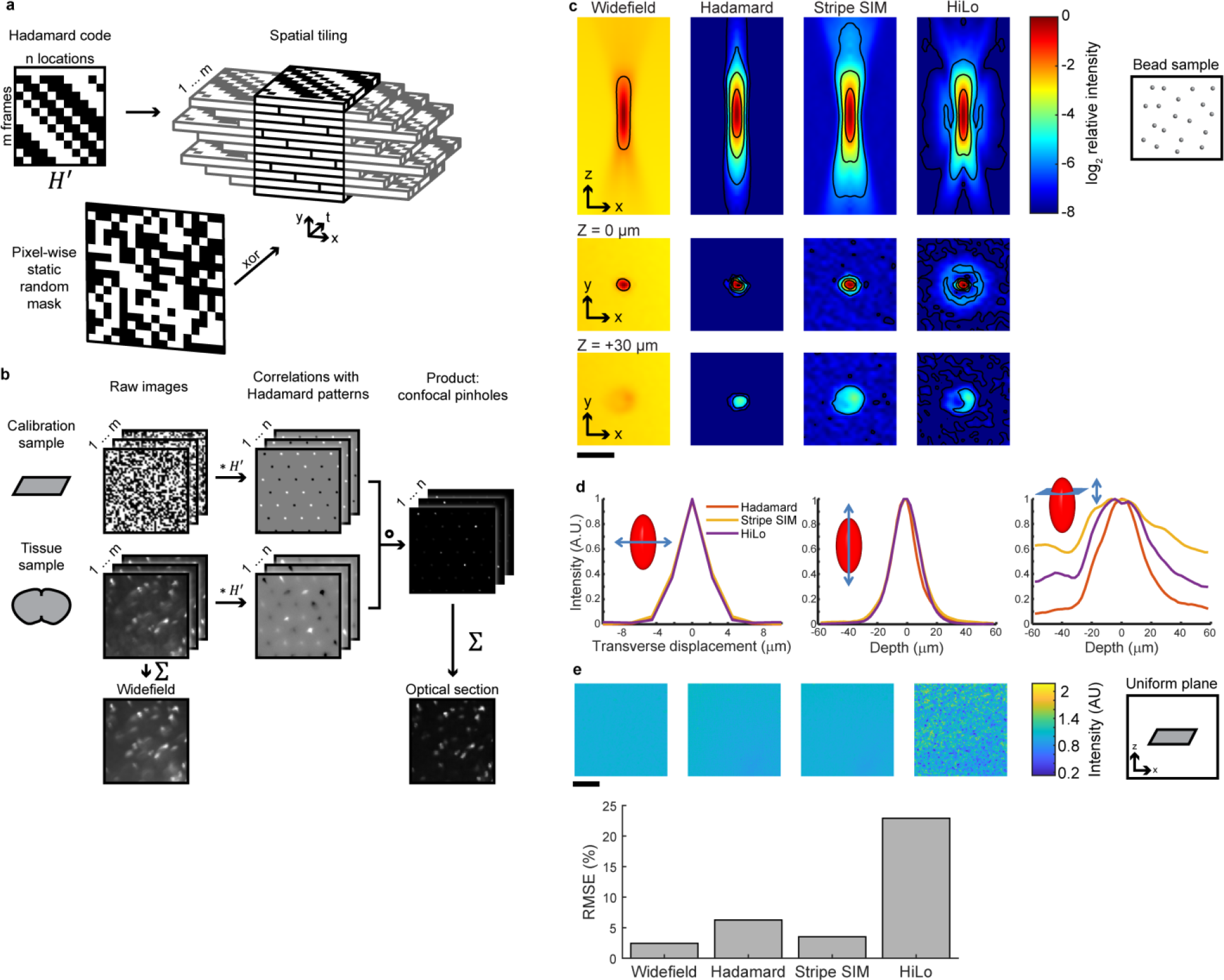
Hadamard microscopy achieves optical sectioning by matched filtering demodulation. (**a**) Codes from a Hadamard matrix were tiled to fill image space. The number of elements in the Hadamard code determined the number of frames in the pattern sequence. A random mask was applied to invert the code in 50% of illumination pixels, yielding pseudorandom patterns with flat spatial and temporal power spectra. (**b**) Raw images were acquired in a calibration sample (a thin homogeneous fluorescent film) and a tissue sample, one frame per Hadamard pattern. Cross-correlation maps between microscope data and Hadamard codes produced arrays of peaks corresponding to signals from distinct sample regions. Negative peaks corresponded to pixels whose Hadamard sequence was inverted. Pixel-wise multiplication of the demodulated images from the calibration sample and from the tissue sample led to multi-point confocal images. These images were summed to produce an image reconstruction. Detailed description in **Methods**. (**c-d**) Characterization of SIM optical sectioning methods using sub-diffraction beads. (**c**) Images show (left to right): Wide-field epifluorescence, Hadamard microscopy using 12 patterns, stripe SIM with period 4 pixels and four phases, and HiLo microscopy using DMD-projected pseudorandom patterns. Top row: Radially averaged meridional cross-section of the point-spread function (PSF). Second row: transverse cross-section at the focal plane. Third row: transverse cross-section at 30 μm defocus. For all (c) the color scale is logarithmic, and contours were drawn on every 4-fold change in intensity. Scale bar is 20 μm. (**d**) (Left and center) Lateral and axial line profiles through the PSF show equivalent resolution for the three sectioning methods. (Right) integrated intensity in transverse cross-sections reveals off-axial spurious side lobes in stripe SIM and HiLo which contribute to out-of-focus crosstalk. (**e**) A uniform fluorescent plane at the focal plane resulted in larger inhomogeneities when imaged using HiLo in comparison with the other methods. The fractional noise in HiLo did not decrease with increasing photon counts. Top: Optical section images of the uniform plane, all shown at the same linear color scale. Bottom: Deviations from uniformity in the images on top. This artifact in HiLo is a known consequence of statistical fluctuations in the total illumination dose, and is not corrected by increasing the exposure time ^41^. Hadamard and stripe SIM microscopies avoided this artifact by providing illumination whose time-average intensity was precisely the same at all sample points. Scale bar is 200 μm. In (c) and (e), each sample was imaged in matched conditions for all methods whenever possible (number of images, illumination intensity, acquisition time). Detailed description in **Methods**.

**Supplementary Figure 3:**
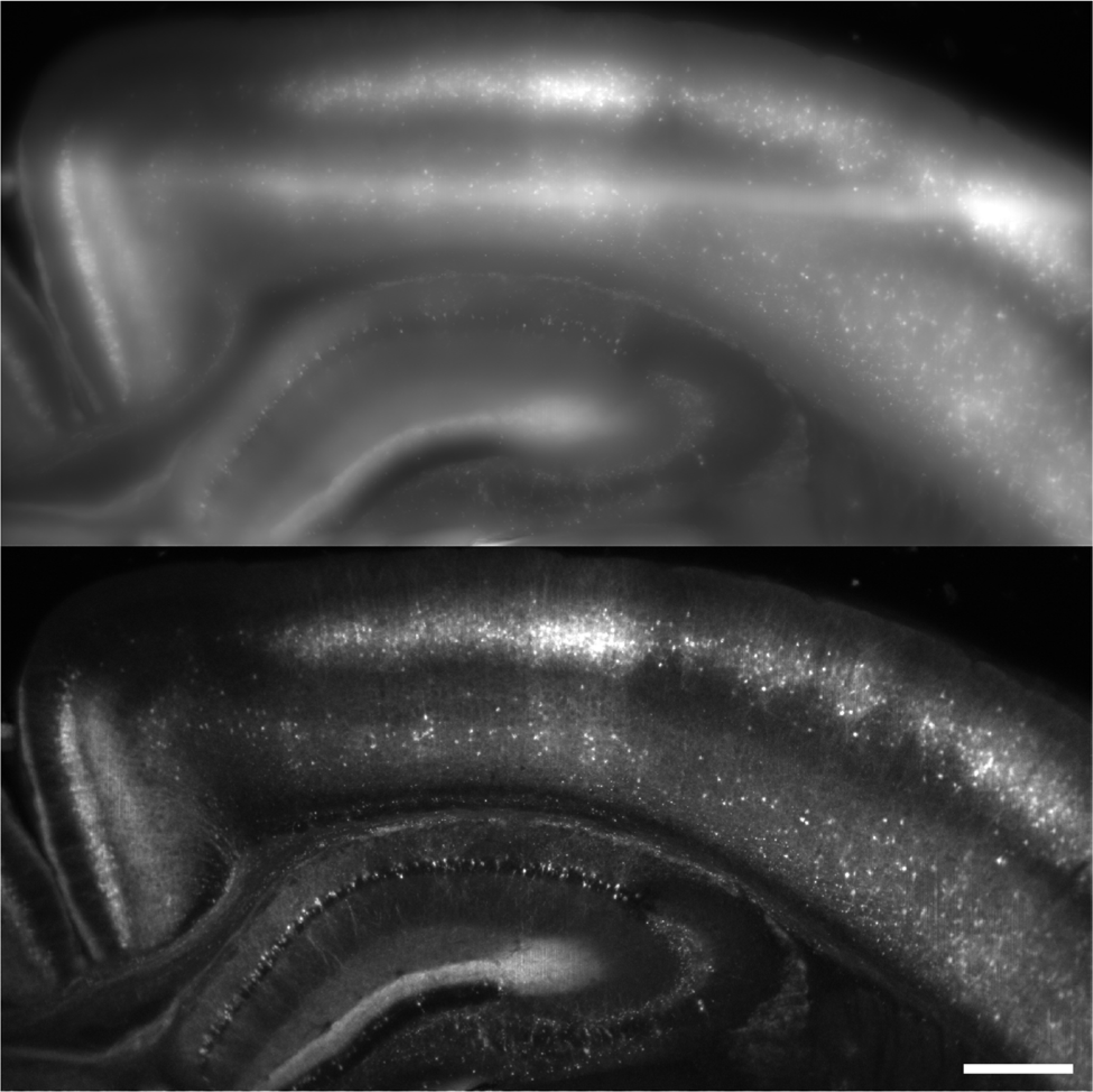
Hadamard microscopy enables wide-field optical sectioning in brain tissue. Maximum-intensity projection of an acute brain slice expressing membrane-targeted Citrine in CaMK2a-Cre positive cells. Top: wide-field epifluorescence. Bottom: Hadamard microscopy. Scale bar 500 μm.

**Supplementary Figure 4:**
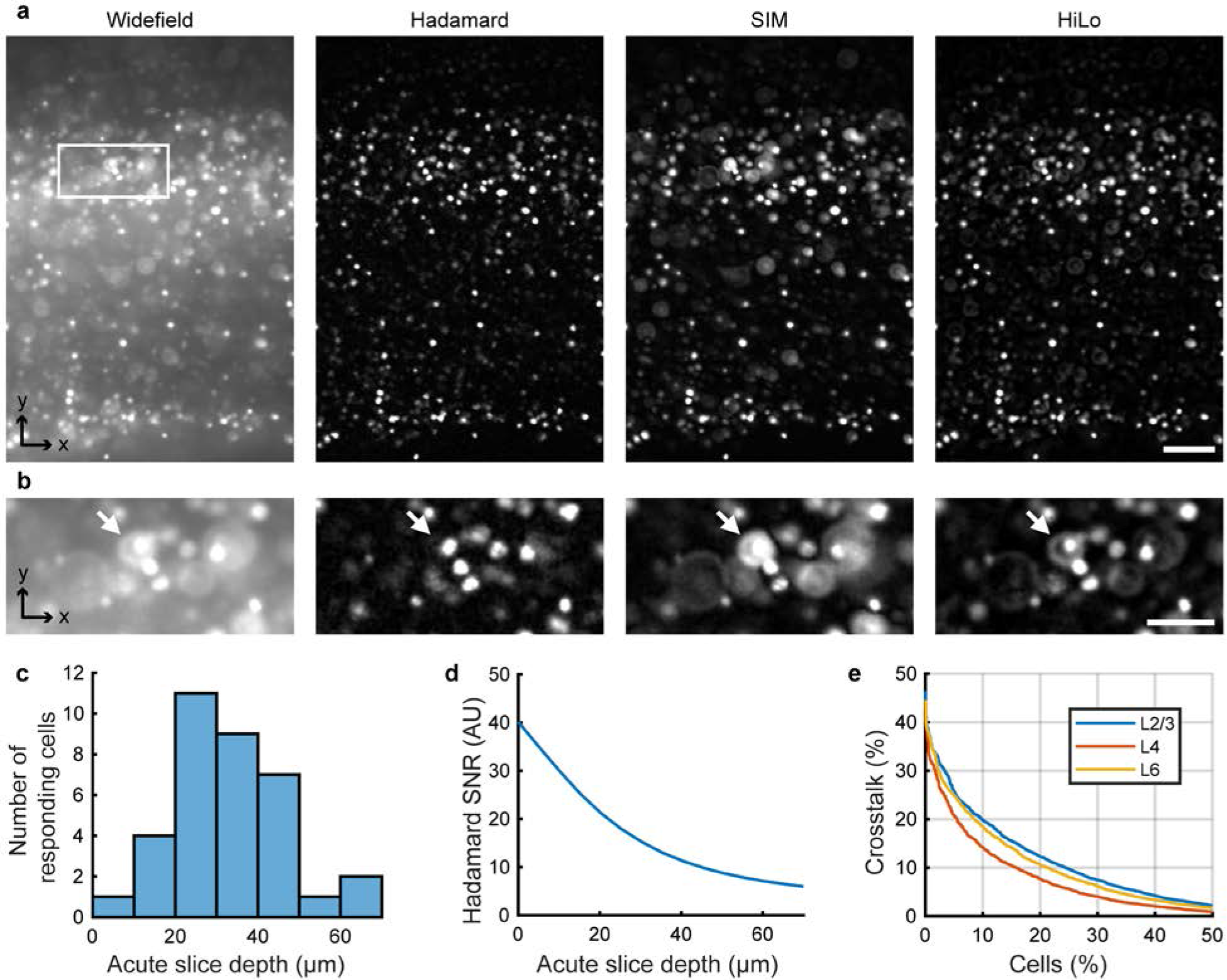
Hadamard microscopy resolves individual H2B-jRGECO1a labeled neurons in acute brain slice. (**a**) Images acquired with different 1P computational optical sectioning methods. Images were acquired in the same sample with matched conditions (number of images, illumination intensity, acquisition time). The sample comprised an acute brain slice expressing H2B-jRGECO1a. Images show (left to right): Wide-field epifluorescence, Hadamard microscopy using 12 patterns, SIM with period 4 pixels and four phases, and HiLo microscopy using DMD-projected speckle patterns. Scale bar 100 μm. The white box region is expanded in (b). (**b**) Hadamard microscopy avoids defocus lobes present using other methods. The white arrows indicate a defocused cell that is rejected by Hadamard microscopy but appears in the other techniques. All images use the same linear scale of normalized grey values. Scale bar 50 μm. (**c**) Depth distribution of responsive cells during Hadamard functional recording, measured by high resolution confocal microscopy acquired after the functional measurement and registered to the Hadamard images. The depth was 32.2 ± 12.7 μm (mean ± std. dev., *n* = 35 neurons). (**d**) Depth-dependent decay in SNR for Hadamard microscopy in acute slices. Decay length was σ_*z*_ = 27 μm. (**e**) Estimated distribution of crosstalk in neuronal recordings using Hadamard microscopy. Only 10% of cells had more than 20% crosstalk in L2/3 (fluorescence attributable to other cells; **Methods**).

**Supplementary Figure 5:**
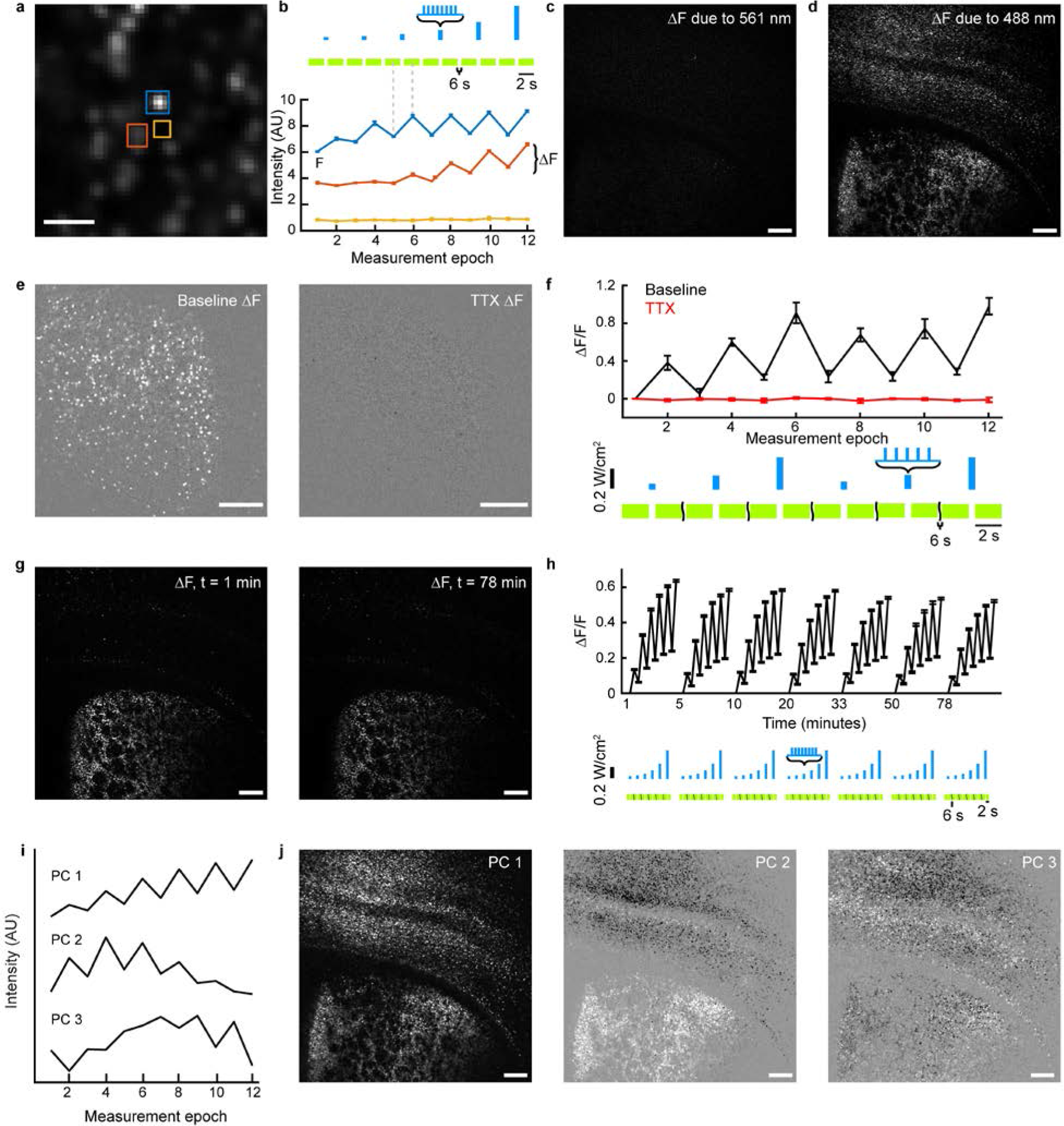
Ultra-widefield AON in acute brain slices. (**a**) Magnified view of region of Fig. 3e showing single-cell resolution. (**b**) Fluorescence traces from regions indicated in (a). Two cells showed optogenetically induced fluorescence transients, while a region between the cells showed no signal. Here the sets of 11 images acquired before and after each optogenetic stimulus were averaged to form single pre- and post-stimulus fluorescence values. Error bars represent s.e.m. over *n* = 11 Hadamard images. Scale bar 25 μm. *F* is defined as the average intensity of the first imaging epoch and Δ*F* is the signal increase following blue light stimulation. (**c**) Measurement of spurious eTsChR activation by yellow (561 nm) light. The image shows the difference between mean fluorescence of H2B-jRGECO1a in the 2^nd^ and 1^st^ second after onset of yellow light for Ca^2+^ imaging. Image represents a mean of *n* = 3 repetitions of the measurement. Spurious eTsChR activation would cause neural firing, which would lead to an increase in H2B-jRGECO1a fluorescence. (**d**) Mean Δ*F* induced by blue light stimulation, averaged over three runs. (c) and (d) are scaled identically. (**e**) Mean Δ*F* images from striatum before (left) and after (right) addition of TTX (1 μM). Images are scaled identically. (**f**) Mean Δ*F*/*F* per measurement epoch from *n* = 360 cells in (f) before TTX addition (black) and after TTX addition (red). Blue light stimulation consisted of 5 pulses at 12.5 Hz of 488 nm light at 60, 120, and 300 mW/cm^2^, repeated twice. (**g**) One slice was repeatedly stimulated and imaged over 78 minutes with protocol in Fig. 3. Mean Δ*F* images from first run (left) and last run (right), scaled identically. (**h**) Average Δ*F*/*F* per measurement epoch for *n* = 3,195 cells in each run in slice shown in (g). (**i**) Waveforms of main principal components from *n* = 31,754 cells. (**j**) Principal components from (i) projected into pixel space for slice in Fig. 3e. Unless otherwise stated, all scale bars 250 μm. Error bars indicate +/-s.e.m..

**Supplementary Figure 6:**
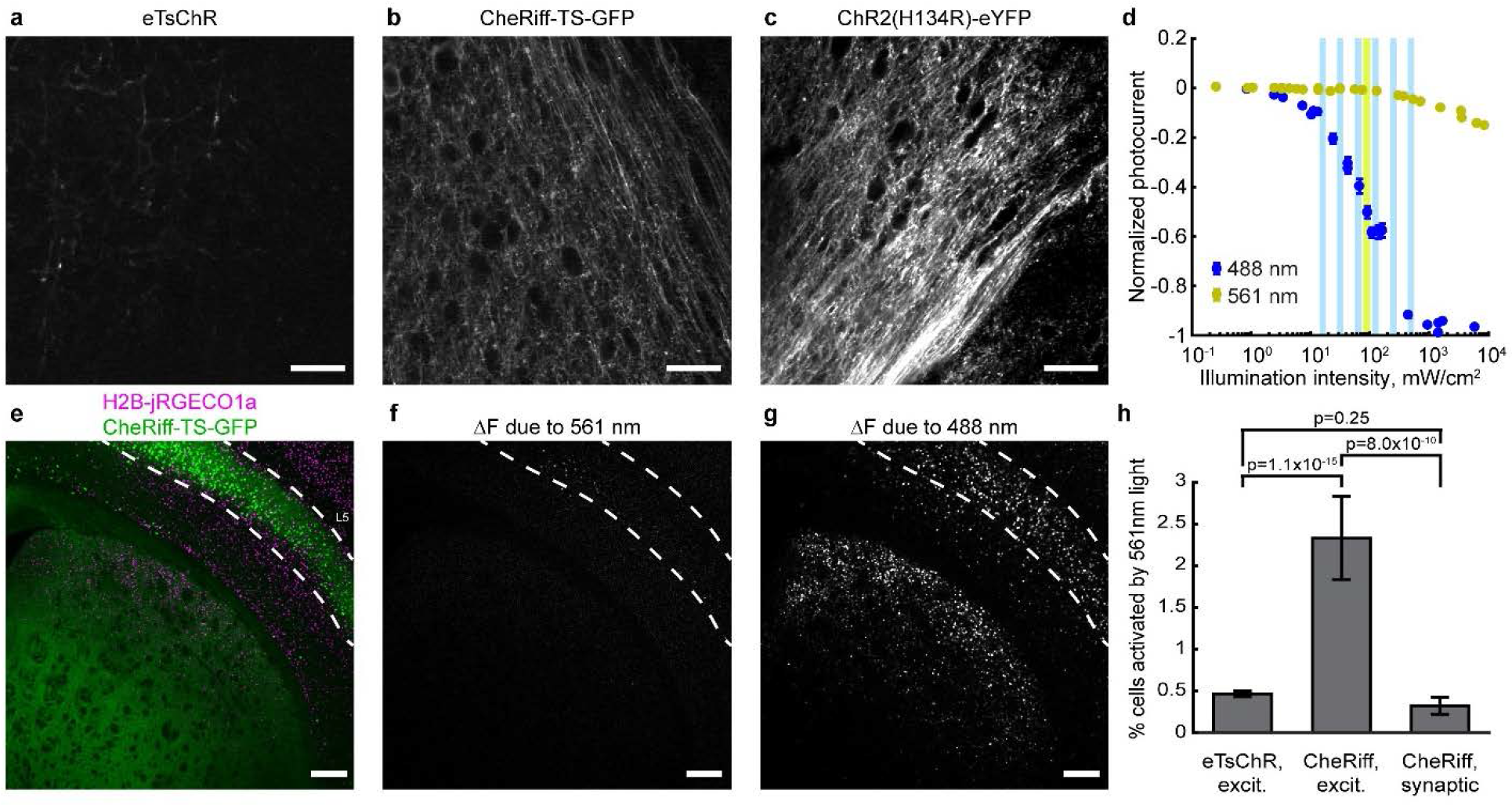
Trafficking of optogenetic actuators in axons in tissue. (**a-c**) Images of axonal trafficking of eTsChR, CheRiff-TS-GFP, and ChR2(H134R)-YFP, scaled to the same counts. Equal volumes of AAV2/9-hSyn-ChR2(H134R)-eYFP, AAV2/9-hSyn-CheRiff-TS-GFP, and AAV2/9-hSyn-eTsChR were injected in the left hemisphere in separate mice and coronal slices of the contralateral hemisphere were prepared after > 4 weeks. Images were acquired near the corpus callosum with 2-photon microscopy. (**d**) Comparison of CheRiff photocurrents in HEK293T cells induced by yellow (561 nm) and blue (488 nm) light. Vertical bars indicate intensities used in acute slice experiments. The blue illumination intensity to achieve 50% activation was 94 mW/cm^2^ (88, 99 mW/cm^2^ 95% confidence interval, n = 7 HEK cells). (**e**) A brain slice expressing CheRiff-TS-GFP (green) in Rbp4-Cre^+^ L5 pyramidal cells and H2B-jRGECO1a (magenta) throughout cortex and striatum was prepared as in Fig. 5a. (**f**) CheRiff activation by yellow (561 nm, 100 mW/cm^2^) light. The image shows the difference between mean fluorescence of H2B-jRGECO1a in the 2^nd^ and 1^st^ second after onset of yellow light for Ca^2+^ imaging. Image represents a mean of *n* = 3 repetitions of the measurement. Spurious CheRiff activation would cause neural firing, which would lead to an increase in H2B-jRGECO1a fluorescence. (**g**) Mean Δ*F* induced by blue light stimulation, averaged over three runs. (g) and (h) are scaled identically. Scale bars 50 μm in (a-d), 250 μm in (e-h). Dashed lines in (f-h) indicate Layer 5 of the cortex. (**h**) Comparison of optical crosstalk for different optogenetic actuators and protocols, as measured by percent of cells showing Ca^2+^ transients in response to onset of illumination with 561 nm light for fluorescence imaging. The three conditions corresponded to eTsChR in the excitability assay (co-expression of actuator and reporter in the same neurons), CheRiff in the excitability assay, and CheRiff in the functional connectivity assay (mutually exclusive expression of actuator and reporter).

**Supplementary Table 1:** *In vitro* characterization of RGECIs. Quantification of action potential responses in cultured neurons in Fig. 2b and Supplementary Fig. 2a, and photobleaching kinetics in HEK293T cells. Action potential magnitudes and sensor kinetics are from 3 FOVs for R-CaMP2 and 4 FOVs for jRGECO1a and jRCaMP1a in separate dishes. Dishes were stimulated with 1 ms field stimulation pulses while imaging RGECI fluorescence at 50 Hz with 2.45 W/cm^2^ 561 nm illumination. Photobleaching measurements were performed in HEK293T cells under 44 W/cm^2^ 561 nm illumination (compared to 0.1 W/cm^2^ used in slice imaging). All values are reported as mean ± s.e.m..

| Construct | Single AP max $\Delta F/F$ (%) | $\tau_{on}$ (ms) | $\tau_{off}$ (ms) | $\tau_{bleach}$ (s) |
| --- | --- | --- | --- | --- |
| jRGECO1a | 54 ± 10, n = 4 FOV, ~30 cells/FOV | 47.2 ± 1.0 | 443 ± 38 | 80.5 ± 5.1, n = 9 cells |
| R-CaMP2 | 31 ± 3, n = 3 FOV, ~30 cells/FOV | 26.3 ± 1.0 | 271 ± 20 | 61.9 ± 2.8, n = 8 cells |
| jRCaMP1a | 17 ± 4, n = 4 FOV, ~30 cells/FOV | 61.2 ± 2.1 | 1600 ± 160 | 37.8 ± 2.1, n = 8 cells |

**Supplementary Table 2:** Patch characterization of eTsChR. Patch parameters of cells in Fig. 2d and Supplementary Fig. 2c-f. All values are reported as mean ± s.e.m., n = 6 cells throughout.

|  | eTsChR | ChR2(H134R)-GFP | *p*-value, Student's *t*-test |
| --- | --- | --- | --- |
| Access resistance (MΩ) | 12.3 ± 1.5 | 12.4 ± 1.3 | 0.96 |
| Membrane resistance (MΩ) | 633 ± 84 | 467 ± 88 | 0.20 |
| Membrane capacitance (pF) | 36.5 ± 4.8 | 44.9 ± 9.7 | 0.45 |
| Resting potential (mV) | -36.5 ± 4.8 | -44.3 ± 2.9 | 0.13 |

## Supplementary Discussion

### Signal-to-noise ratio of Hadamard microscopy

Here we compare the signal-to-noise ratio (SNR) properties of multi-focal scanning confocal vs. Hadamard microscopy, both implemented in the same DMD-based system, limited in total illumination time and maximum intensity. In multi-focal scanning confocal, isolated pixels on the DMD are illuminated sequentially in a tiled array, and synchronized images are acquired on the camera. A computational pinhole is imposed by only retaining signal from the camera pixels matched to the currently active DMD pixels.

For simplicity, let us assume that there is a 1:1 mapping of DMD pixels onto camera pixels. Let us tile the image into blocks of *N* pixels. Let *S* be the mean number of signal photons acquired in one camera pixel in one frame when the corresponding DMD pixel is turned on. Let α be the mean number of background photons acquired in camera pixel *i* in one frame when DMD pixel *j* ≠ *i* is turned on, averaged over all *j* within a block of *N* pixels.

Both Hadamard and multi-focal scanning require acquisition of *N* camera frames to assemble one composite image. Let us assume that *N* is large enough that we can ignore crosstalk of illumination between blocks.

The shot-noise-limited signal-to-noise ratio is:

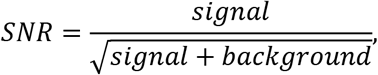

where all intensities are measured in photon counts.

### Multi-focal confocal

In a single composite image of *N* frames, each pixel is illuminated once, so the signal level is *S*. There is no background. The shot noise-limited signal-to-noise ratio is 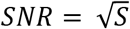.

### Hadamard

In a single composite image of *N* frames, each pixel is illuminated *N*/2 times, so the signal level is *S N*/2. The background level is α(*N* − 1)*N*/2 (there are *N* – 1 pixels that contribute to background, each is illuminated for *N*/2 frames), where α is the mean pixel-to-pixel crosstalk. In practical use, *N* ≫ 1 (*N* = 12 in our functional measurements and *N* = 64 in structural measurements), so the *SNR* is approximately:

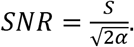

If α < *S*/2, then Hadamard wins; otherwise multi-focal confocal wins.

We find experimentally that for *N* = 12, α/S = 0.51, indicating comparable shot-noise for the multi-focal and Hadamard approaches. For *N* = 64, α/S = 0.26, indicating superior performance of Hadamard over multi-focal confocal.

## Materials and methods

- DNA constructs
- Cell culture and gene expression

○ HEK cells culture and gene expression
○ Low titer lentivirus production
○ Primary neuron culture and gene expression
- Imaging and electrophysiology in culture

○ Microscope
○ Imaging and electrical recordings
○ Data analysis
- Hadamard Imaging

○ Microscope
○ Illumination patterns
○ Calibration
○ Reconstruction
○ Hadamard image formation
○ Image processing and filtering
○ Characterization
- Animals and slice measurements

○ Animals used
○ AAV injection
○ Acute slice preparation
- Analysis of slice data

○ Registration
○ Cell selection
○ Exclusion of spontaneously active and dying cells.
○ Generation of excitability maps.
○ Cortical layer analysis.

### DNA constructs

R-CaMP2 was a gift from Haruhiko Bito. TsChR was a gift from Ed Boyden. jRGECO1a and jRCAMP1a were obtained from Addgene (Plasmids #61563 and #61562). All RGECIs were cloned between the BamHI and EcoRI sites of the backbone from FCK-Arch-GFP (Addgene Plasmid #22217) for expression in cultured neurons and for lentiviral production. For photophysical characterization, RGECIs were also cloned into an analog of the FCK vector replacing the CaMKIIα promoter with a CAG promoter, a configuration we refer to as FCAG. The jRCaMP1a and jRGECO1a constructs included the nuclear export sequences found in the original publication^16^. For nuclear localization, the nuclear export sequence of jRGECO1a was replaced with an H2B tag, and cloned into an AAV-hSyn-DO Cre-off vector. TsChR, including an N-terminal Kir2.1 trafficking sequence followed by a GFP fluorescent tag, was cloned into FCK and into an AAV expression vector under control of the human synapsin promoter (AAV-hSyn). CheRiff-TS-GFP (Addgene Plasmid # 51693) was cloned into an AAV-CAG-DIO expression vector. FCK-ChR2(H134R)-GFP was used as a reference for eTsChR characterization. FCK-VSV-G (Addgene Plasmid #8454) and psPAX2 (Addgene Plasmid #12260) were used in lentiviral production. pUC19 (NEB #N3041) was used as a diluent in calcium phosphate transfections.

### Cell culture and gene expression

#### HEK cell culture and gene expression

Photophysical measurements of RGECIs were performed in HEK293T cells (ATCC CRL-11268) cultured as previously described^13^. Cells were grown at 37 °C, 5% CO_2_ in DMEM containing 10% FBS (Life Technologies 10082-147) and 50 U/mL penicillin-streptomycin (Life Technologies 15060-063). Cells were split with trypsin-EDTA (Life Technologies 25300054) every 2-3 days and used before passage 25. For gene delivery, cells were grown to 70% confluence in 24 well plates or 35 mm plastic dishes. 200 ng (for 24 well plates) or 400 ng (for 35 mm plastic dishes) of FCAG-RGECI DNA was transfected using TransIT-293 (Mirus 2705) following manufacturer instructions. After 24 hours, cells were split onto Matrigel (Fisher Scientific 356234) coated glass bottom plates (In Vitro Scientific D35-14-1.5-N) and imaged 24 hours later.

#### Low titer lentivirus production

HEK293T cells were cultured as in the previous section, except that cells were split daily and the cell density was always maintained between 30 and 70%. Prior to P11, cells were split onto gelatin coated plates, prepared by incubating 15 cm plastic dishes (Nunc) for 20 minutes at room temperature with 10 mL EmbryoMax 0.1% Gelatin solution (Millipore FS-006-B) and aspirating to dryness. 10 cm dishes were also used, and all amounts were scaled to the smaller surface area. After cells reached 80% confluency, cells were switched to 16 mL pre-warmed DMEM without FBS for 1-2 hours. For each dish, the following were added, in order, to 1.2 mL DMEM: 14 μg of FCK-RGECI plasmid, 9 μg psPAX2, and 4 μg VsVg were combined with 36 μL of 1 mg/mL PEI in water (Aldrich #408727). The tube was vortexed and incubated at room temperature for 10 minutes. The mixture was then pipetted dropwise over the surface area of the dish and the cells were returned to the incubator for 4 hours. After the incubation, the medium was replaced with 16 mL DMEM + 10% FBS without antibiotics. 36-48 hours later, the medium was collected and centrifuged for 5 min at 500 × g. The supernatant was filtered through a 0.45 μm filter blocked with DMEM + 10% FBS and aliquoted in 1-5 mL fractions. Aliquots were kept at -80°C until use.

#### Primary neuron culture and gene expression

Cultured rat hippocampal neurons on astrocyte monolayers were prepared as previously described^13^, with two modifications: (1) In Vitro Scientific dishes model D35-14-1.5-N were used instead of D35-20-1.5-N, while keeping the cell densities the same and (2) neurons were cultured in Neurobasal-A (Life Technologies 10888-022) supplemented with B27 (Life Technologies 17504044) instead of Brainbits’ NbActiv4. For electrophysiological and AON measurements, neurons were transfected via calcium phosphate, as previously described^13^ on DIV7 and used on DIV14-16. For comparison of RGECI performance by field stimulation (**Supplementary Fig. 1a-b**), cultured neurons were lentivirally transduced. On DIV 7, half of the media from each dish (1 mL) was reserved and replaced with 250 μL of low titer FCK-RGECI lentivirus. After two days, all of the media was removed and replaced with the reserved media supplemented with an additional 1 mL of Neurobasal-A + B27 supplement.

### Imaging and electrophysiology in culture

#### Microscope

A custom-built epifluorescence microscope was used for measurements in HEK293T cells and in cultured neurons. Illumination was provided by a 561 nm 100 mW laser (Cobolt Jive 0561-04-01-0100-500) or a 488 nm 100 mW laser (Coherent Obis 1226419). The laser lines were combined and focused in the back focal plane of the objective (Olympus Fluor 4x 0.24 NA for single action potential measurements of RGECIs; Olympus LCPlanFL 20x 0.40 NA for RGECI photobleaching measurements; Olympus UPlanSApo 10x 0.40 NA for RGECI photoswitching characterization; Olympus ApoN 60x 1.49 NA Oil for eTsChR characterization). Fast modulation of the 488 nm laser was achieved with an acousto-optic tunable filter (Gooch&Housego TF525-250-6-4-GH18A). Both laser lines were additionally modulated by neutral density filters as necessary. Fluorescence light was separated from illumination light using a quadband dichroic (Semrock Di01-R405/488/561/635). HQ550/50m or ET595/50 bandpass emission filters (Chroma) were used to isolate GFP or RGECI fluorescence, respectively, before capturing on a scientific CMOS camera (Hamamatsu Orca Flash 4.0). For photobleaching measurements, an additional 1 OD filter was inserted in the imaging path to avoid saturating the camera. Illumination profiles were acquired on bead samples before experiments each day and spot size was determined using a 1/e^2^ cutoff. Laser powers were measured at the sample plane. A digital acquisition (DAQ) card (National Instruments PCIe 6259) was used to synchronize command and recording waveforms. Imaging frame rates and illumination powers are indicated in figure captions for each experiment.

#### Imaging and electrical recordings

In all imaging measurements, culture medium was replaced with imaging buffer containing, in mM, 125 NaCl, 2.5 KCl, 2.5 HEPES, 30 glucose, 1 MgCl_2_, 3 CaCl_2_. The buffer pH was adjusted to 7.3 and osmolarity was 310 mOsm. Measurements were carried out at room temperature. 10 μM CNQX, 20 μM gabazine, and 25 μM APV (all Tocris) were included in cultured neuron experiments to block synaptic transmission. Channelrhodopsin characterization measurements were performed in synaptic blockers with the addition of 1 μM tetrodotoxin (Tocris). No additional all-*trans* retinal was added.

Field stimulation (**Supplementary Fig. 1a-b**) was performed by inserting two chlorided silver wire loops 2 cm apart into the glass-bottomed imaging dish, touching the plastic on either side of the coverslip. A high voltage amplifier (Krohn-hite 7600M) was used to amplify 1 ms pulses generated by the DAQ card to 60-120 V. 3-4 FOVs were acquired for each construct, using a fresh dish each time.

For patch clamp electrophysiology measurements (**Fig. 1c, g**, **Supplementary Fig. 1c-d, Supplementary Fig. 6d**), 3-5 MΩ borosilicate glass pipettes (WPI) were filled with internal solution containing, in mM, 125 potassium gluconate, 8 NaCl, 0.6 MgCl_2_, 0.1 CaCl_2_, 1 EGTA, 10 HEPES, 4 Mg-ATP, 0.4 Na-GTP, adjusted to pH 7.5 and 295 mOsm with sucrose. Voltage- and current-clamp recordings were obtained with a Multiclamp 700B amplifier (Molecular Devices) while illuminating with 1 s 488 nm pulses or 2s 561 nm pulses of intensities indicated in figure captions. In voltage clamp measurements, cells were held at -65 mV. In current-clamp measurements, an offset current was injected to maintain the resting membrane potential at -65 mV. Signals were filtered at 5 kHz with the amplifier’s internal Bessel filter and digitized at 10 kHz.

#### Data analysis

All values are expressed as mean ± standard error of the mean (s.e.m.). *P* values were obtained from Student’s *t*-tests unless otherwise indicated.

Whole FOV RGECI single action potential responses (**Supplementary Fig. 1a-b, Supplementary Table 1**) were extracted as previously described^42^. Activation time constants were extracted from monoexponential fits between stimulation onset and maximum ΔF/F. For inactivation time constants, the fluorescence trace after the maximum ΔF/F was fit to a sum of two exponential decays, and the τoff was taken as the time for the fit to decay to half its maximum value. Photobleaching traces (**Supplementary Table 1**) were extracted from separate cells and fit to a monoexponential to obtain time constant τ_bleach_.

Movies of blue light photoswitching (**Fig. 1f**, **Supplementary Fig. 1g-h**) were preprocessed to reject saturated pixels and a threshold equal to half the average of movie was used to separate foreground from background. Background intensity was subtracted from the original movies and the averages of the resulting foreground traces (combining 10-20 cells each) were used in downstream analysis. Traces were converted to ΔF/F using the fluorescence value before blue light stimulation as F_0_. “Photoswitching ΔF/F” was defined as the ΔF/F immediately after blue light illumination ends (**Fig. 1f**, inset).

For comparison of channelrhodopsins (**Fig. 2d**, **Supplementary Fig. 3c-e**), cells were rejected if they required >100 pA holding current to maintain -65 mV in current clamp or if their baselines drifted by more than the smallest steady state photocurrent amplitude in voltage clamp mode. Steady-state 488 nm photocurrents were extracted as the average photocurrent over the last 100 ms of blue light illumination. Steady state 561 nm photocurrents and depolarizations were extracted from 1 s of data. On time constants were obtained from single exponential fits to the first 1.5 ms of 488 nm illumination. Off time constants were obtained from single exponential fits to the 99.5 ms following blue light 488 illumination.

Recordings of jRGECO1a fluorescence in **Fig. 1d** were corrected for photobleaching with a di-exponential fit to the initial period in each movie, before stimulation, while recordings of BeRST1 fluorescence were corrected for photobleaching by a sliding, 1000 point, median filter. Both traces were converted to ΔF/F based on the fluorescence before blue light stimulation. Frames acquired during blue light stimulation were dropped to avoid optical crosstalk.

### Hadamard imaging

#### Microscope

In the ultra-widefield microscope (**Fig. 2a**), a 561 nm laser beam (MPB Communications F-04306-02) was transmitted through a rotating diffuser, and merged with a 470 nm LED beam (Thorlabs M470L3). Both were expanded, focused, and coupled through free space to fill with high NA illumination a digital micromirror device (DMD) module (Vialux V-7001; 1024×768 pixels, 13.7 μm pitch). Multiple diffraction orders emitted from the DMD pattern were transmitted by a 100 mm projection tube lens (Zeiss Makro-Planar 100 mm, L1 in **Fig. 2a**), reflected off a custom dichroic mirror (Semrock Di01-R405/488/561/635-t3-60×85), and imaged onto the sample by a 50 mm objective lens (Olympus MVPLAPO 2XC, NA 0.5, L2 in **Fig. 2a**). The 3 mm substrate thickness of the dichroic mirror minimized warping-induced projection aberrations. Fluorescence emission was collected through the same objective and dichroic, a large diameter (60mm) emission filter (Semrock FF01-520/35-60-D or Chroma ET600/50m, F in **Fig. 2a**), and a 135 mm imaging tube lens (Zeiss Apo-Sonnar 135 mm, L3 in **Fig. 2a**) onto a scientific CMOS camera (Hamamatsu Orca Flash 4.0, 2048×2048 pixels). The FOV was 4.6×4.6 mm^2^ in the sample plane, corresponding to a magnification of 2.89x onto the camera, and 2.17x onto the DMD. Camera and DMD pixels were 2.25 μm and 6.3 μm wide in the sample, respectively. Hardware and triggers were programmed in LabView, with pattern generation and data analysis performed in MATLAB.

#### Illumination patterns

To reject light scattered within the sample, pattern sequences were designed such that in the projected series of 2D images, neighboring locations of the sample were illuminated with orthogonal functions of intensity vs. time. A Hadamard matrix, *H*, of size *m* is a binary square matrix with elements {−1,1} that fulfills *H*^*T*^*H* = *mI*_*m*_, where *I*_*m*_ is the identity matrix of size *m*; its normalized form has value 1 in the first column and first row. Illumination intensities could not be negative, so the projected intensity patterns were defined as *P* = (*H*′ + 1)/2 where *H*′ = *H*[1, …, *m*; *m* − *n* + 1, …, *m*] was an incomplete orthogonal basis given by the last *n* columns of a normalized Hadamard matrix, with *n* < *m*.

The illumination patterns *P* thus had binary values {0,1} corresponding to DMD mirror positions OFF and ON respectively. Each location was illuminated with a positive temporal function orthogonal to all other designed Hadamard codes, as verified by *P*^*T*^*H*′ = *I*_*n*_ * *m*/2. For a given number of locations, a Hadamard matrix provided a set of shortest possible binary orthogonal functions. To arrange the *n* codes in *P* into illumination patterns, *m* = *n* + 1 images were defined assigning code *k*_*i,j*_ ∈ {1.. *n*} to DMD pixel (*i*, *j*), as *k*_*i,j*_ = mod(*i* * *q* + *j*, *n*) + 1, where *q* was an offset parameter that maximized spatial separation of repeated codes. (*n*, *q*) was set to (11,3) for functional imaging, and to (63,14) or (59,8) for structural imaging. To further reduce spurious scattering cross-talk, a random binary mask *R* was generated to flip the sign of 50% of DMD pixels, applied as an exclusive OR operation on all DMD patterns against the same mask *R*.

#### Calibration

To prepare the system for each imaging session, a calibration data-set *C* was obtained by placing a thin fluorescent sample at the focal plane, and acquiring an image with each illumination pattern. The sample consisted of green or orange neon Sharpie (Newell Brands, NJ) ink painted on (or sandwiched between) glass coverslips, to match imaging conditions of subsequent acute (fixed) tissue experiments. For each camera pixel, the time series of its photon counts was cross-correlated against each Hadamard sequence as *C^T^H*′. The resulting cross-correlation images displayed sharp peaks indicating the projected DMD locations for each code, with positive or negative correlation given by *R*. A synthetic approximation to the cross-correlation maps was calculated by finding the code with maximum absolute correlation for each pixel, yielding homogeneous, noise-free cross-correlation maps.

#### Reconstruction

A Hadamard sequence data-set *D* was acquired after replacing the calibration sample with a tissue sample. Photon counts at each camera pixel were cross-correlated against each Hadamard sequence as *D*^*T*^*H*′. Cross-correlation images displayed a set of peaks modulated by the local fluorophore density, and broadened by off-focus fluorescence and light scattering in the sample. Each peak characterized the scattering function of the corresponding tissue location, i.e. its absolute value represents the image one would record with an illumination spot focused solely at that location in the tissue. The next step was to apply a set of computational ‘pinholes’ to select the unscattered in-focus photons, and to reject all others. The spatial filter was implemented through the element-wise product of calibration correlation maps and tissue correlation maps, resulting in the positive filtered maps *F* = *C*^*T*^*H*′ ∘ *D*^*T*^*H*′. This computational process was akin to sifting emitted light through an array of pinholes as happens physically in spinning disk confocal microscopy. The final computation step was to aggregate the unscattered light by direct sum of the filtered images over all code maps, defining an optical section image 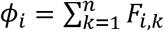.

All image computations in this work were accelerated by computing 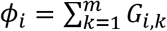, with *G* = *C*^*T*^ ∘ *D*^*T*^. This approach is numerically equivalent to the more involved process described above, as proved by:

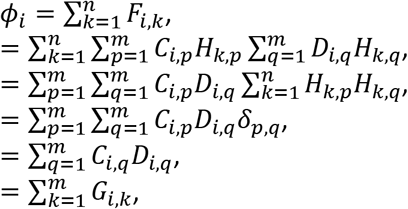

where δ_*p,q*_ is a Kronecker delta. The resulting optical section preserved unscattered light emitted from the focal plane, while rejecting scattered light and background emissions. Standard wide-field epifluorescence images were also computed from each Hadamard dataset by computing a direct sum of all frames in the raw images, 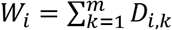.

To correct for slight motion artifacts due to sample drift, all datasets from one brain slice were registered to a reference image using a b-splines transform maximizing mutual information^43^.

#### Hadamard image formation

To understand the optical sectioning process, Hadamard microscopy was modeled as an incoherent illumination, intensity-linear space-invariant optical system, in which the intensity after propagation is given by a convolution between intensity before propagation and an intensity impulse response function. In a discrete representation, the circulant convolution matrix 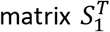 represented three-dimensional excitation intensity at the object, in response to an impulse function reflectance at the DMD plane (turning on one DMD pixel). Similarly, *S*_2_ was defined as the intensity collected by an impulse detector at the camera plane from emitted fluorescence in a three-dimensional object (analogous to detection from one camera pixel). The data collected from tissue with fluorophore distribution *G* upon illumination with a structured illumination pattern *P* was represented as 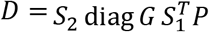, where diag denoted rearrangement between vector and diagonal matrix. Calibration with a thin uniform fluorescent object and no scattering was represented as *C* = *P*. After assuming that *P* contains an orthonormal Hadamard code with no spatial repetition, it followed that *C*^*T*^*H*′ = *I*_*n*_, and 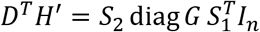. Then 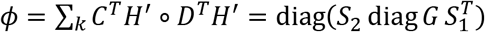, or ϕ = (*S*_1_ ∘ *S*_2_)*G*. The reconstructed optical section ϕ was proportional to the object *G* convolved with the confocal scattering function *S* = *S*_1_ ∘ *S*_2_ that resulted from the element-wise product of the projection and collection scattering functions.

#### Image processing and filtering

The lines between DMD pixels led to a periodic grid artifact in Hadamard optical sections. A Gaussian stopband filter was used to attenuate these artifacts. The filter parameters were not changed after initial set-up.

The size of the computational pinholes could be adjusted in software to trade optical signal level for z-resolution. Tuning of pinhole sizes was achieved by applying a spatial Gaussian filter to the calibration patterns, with σ = 5.6 μm for functional images, and σ = 3.4 μm for structural images. Further increases in σ to sizes larger than the spacing of pinholes resulted in a continuous transition to wide-field epifluorescence imaging.

An additional source of systematic error came from local inhomogeneity of illumination patterns. While the projected patterns have 50% duty cycle on average, variations in local illumination can change the relative contributions of in-plane signal and background, resulting in imperfect background cancellation manifested as regions with periodic background artifacts. This effect was minimized for Hadamard images in **Supplementary Fig. 3** by dividing raw tissue data by its low spatial frequency component, calculated with a Gaussian filter with σ=33.8 μm for m=60, and σ=22.5 μm for m=12. Images in all figures were linearly mapped to grayscale setting 0 to black and saturating to white the 0.01 percentile of highest intensity values unless otherwise indicated.

#### Characterization

We quantified the performance of Hadamard, stripe SIM, and HiLo optical sectioning methods by three measurements. First, we measured the point spread functions by imaging 200 nm fluorescent beads (Invitrogen F8763) embedded in 1.5% agarose gel. Second, we tested the in-plane uniformity of optical sections by measuring a thin fluorescent plane of orange neon Sharpie (Newell Brands, NJ) ink painted on a glass coverslip. Third, we acquired multi-plane images of an acute brain slice expressing H2B-jRGECO1a to evaluate the imaging quality of each method in turbid tissue.

For the beads and plane experiments, illumination patterns for Hadamard codes of length 12, together with striped illumination with period 4 pixels and 4 phases, were interleaved and repeat-averaged to match total photons and photobleaching conditions across datasets. HiLo optical sections were computed from the same patterns used for Hadamard imaging, using a photon-matched uniform illumination image and a repeat-averaged structured image corresponding to one Hadamard pattern. Because HiLo necessarily uses a non-uniform total photon count across the sample, we used more total photons in HiLo optical sections to avoid penalizing this method in the comparison. A series of images taken at Δz = 2.24 μm were acquired to map the three dimensional PSF.

Hadamard images were calculated as 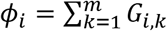, with *G* = *C*^*T*^ ∘ *D*^*T*^. Stripe SIM optical sections were calculated as 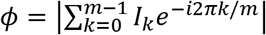, with m=4. HiLo optical sections were calculated setting the wavelet filter σ = 0.75. DMD modulation grid artifacts were present in all datasets and were not corrected. Widefield reference images were obtained by summing all patterns in the Hadamard sequence.

Images of the homogeneous fluorescent plane were acquired following the same protocol as for the beads. The same flat field correction was applied to all datasets by subtracting the offset and dividing by the blurred intensity distribution of a focused widefield image. All datasets were filtered equally to reduce DMD grid artifacts. Within a region of interest, the standard deviation of values was normalized by their mean to obtain coefficients of variation.

### Animals and acute slice measurements

#### Animals

All procedures involving animals were in accordance with the National Institutes of Health Guide for the care and use of laboratory animals and were approved by the Institutional Animal Care and Use Committee (IACUC) at Harvard University. Excitability measurements and characterization of functional Hadamard imaging were performed in wild type C57Bl6 (Charles River Labs #027) mice. Functional connectivity assays were performed in Rbp4-Cre^+/−^ mice donated by Bernardo Sabatini’s lab and originally generated in the GenSat project (#KL100). For structural imaging of membrane bound mCitrine, FLOXed Optopatch-3 mice (Jackson Labs #029679) were crossed with Rbp4-Cre^+/−^ mice or with CaMK2a-Cre^+/−^ mice (Jackson Labs, #005359).

#### AAV injection

AAV2/9-hSyn-DO-H2B-jRGECO1a (1.60×10^13^ GC/mL) and AAV2/9-hSyn-eTsChR (2.22×10^13^ GC/mL) were produced at Massachusetts Eye and Ear Infirmary Vector Core. AAV2/9-CAG-DIO-CheRiff-TS-GFP (5.80×10^13^ GC/mL) was produced by the Stanford Vector Core. AAV1-hSyn-NES-jRGECO1a (2.44×10^13^ GC/mL) was purchased from the University of Pennsylvania Vector Core. When two viruses were coinjected, they were mixed in a one-to-one volume ratio. The final mixture was mixed in a 7:1 ratio with 0.4% Trypan Blue to aid in visualization during injection. For viral injections, neonatal (P0-2) animals were cold-anesthetized and taped to an aluminum heatsink submerged in an ice bath, with their heads resting on a modeling clay support. A stereotaxic injector (WPI #UMC4) mounted on a stereotaxic frame (Stoelting) was used to inject virus 1.6 mm anterior and 1.6 mm lateral to lambda every 0.4 mm starting from 3 mm beneath the surface of the skull. 40 nL of virus was delivered at each depth at a rate of 5 nL/s. If only one virus was used, only 20 nL were injected per depth. Expression levels were sufficiently high for Hadamard imaging from 12 days until at least 9 weeks after injection.

#### Preparation of fixed slices

Fresh 300μm brain sections were incubated in 4% paraformaldehyde overnight at 4 °C, then mounted on a glass slide in Fluoromount and stored at 4 °C.

#### Acute slice preparation and imaging

Acute slices were prepared from P21-28 animals. Animals were deeply anesthetized via isoflurane inhalation and transcardially perfused with ice-cold choline cutting solution, containing, in mM 110 choline chloride, 25 sodium bicarbonate, 2.5 potassium chloride, 7 magnesium chloride, 0.5 calcium chloride, 1.25 monobasic sodium phosphate, 25 glucose, 11.6 ascorbic acid, and 3.1 pyruvic acid (310 mOsm/kg). The brain was blocked with one coronal cut just anterior to the tectum and mounted with Krazy glue on the specimen disk of a Leica VT1200s vibratome. After mounting, hemispheres were separated with a sagittal cut down the midline of the brain. The brain was covered with more ice-cold choline solution and then sliced in 300 μm steps. Slices containing the striatum were recovered for 45 minutes in a 34 °C artificial-cerebrospinal fluid (ACSF) bath containing, in mM, 125 NaCl, 2.5 KCl, 25 NaHCO_3_, 2 CaCl_2_, 1 MgCl_2_, 1.25 NaH_2_PO_4_, 25 glucose (295 mOsm/kg). Slices were kept in room temperature ACSF until ready to measure and were used within 8 hours. All solutions were bubbled with carbogen (95% O_2_, 5% CO_2_) for the duration of the preparation and subsequent experiment.

For imaging, slices were mounted on Poly-L-Lysine (PLL) coated coverslips. Coverslips (Fisher #12-545-80) were plasma cleaned for 3 minutes, covered with 50-100 μL 0.1% (w/v) PLL (150-300 kD) solution (Sigma #P8920) and allowed to dry under vacuum. Coverslips were thoroughly washed with nanopore water and dried before use. To mount the tissue, a slice was transferred to the PLL-coated face of the coverslip with a Pasteur pipette. Excess ACSF was pipetted or wicked away with filter paper, in the process flattening out the brain slice and adhering it to the glass. We found that this method worked reliably for coronal slices from one hemisphere but not for coronal slices from the entire brain. Coverslips were placed in a custom built flow chamber with a microscope slide bottom and #1.5 coverslip lid. ACSF was perfused at a rate of 1 mL/min with a VWR peristaltic pump.

The imaging protocol consisted of a 2 s imaging epoch followed by a 400 ms stimulation period and another 2 s imaging epoch. Each imaging epoch comprised 11 frames of functional Hadamard acquired with a 180 ms period under 100 mW/cm^2^ 561 nm illumination. Blue light stimulation protocols are described in figure captions. The slice was allowed 6 s to recover before starting another imaging epoch. One run consisted of 6 imaging and stimulation rounds over one minute. Runs were repeated several times, spaced out by at least five minutes. NBQX and CPP, or TTX (Tocris) or retigabine, phenytoin, or carbamazepine (Sigma) were added to the ACSF from 1000x stock solutions after several baseline runs.

### Analysis of slice data

#### Registration

After reconstruction of Hadamard images (see Methods above), frames for each epoch were averaged together. Small movements and deformations in the slice over the course of multiple runs were corrected by automatic non-rigid registration^43^. Functional Hadamard recording and structural Hadamard images were manually registered using a 2D affine transformation.

#### Cell selection

ΔF images were calculated for each registered run by subtracting images acquired before blue light stimulation from images acquired immediately after blue light stimulation. Peaks in ΔF images corresponded to individual cells, but noise in ΔF varied as a result of brightness inhomogeneities in the slice, making it difficult to extract peaks directly. To correct for this noise, a widefield image for each slice was blurred with a 2D Gaussian with an 8 pixel (19.2 μm) standard deviation, to remove nucleus sized objects. The square root of this image was used to normalize the ΔF image of the slice. High spatial frequency noise was removed with a 2d Gaussian filter with a 0.5 pixel (1.2 μm) standard deviation. Regions without expression were manually selected and standard deviations in these regions were chosen as a noise floor. Cells were identified as peaks in the normalized ΔF image which had an amplitude larger than the noise floor by a user-defined factor, typically 7 – 10. Cells were required to have a minimum distance in space of 4 pixels (9.6 μm) to avoid double counting cells. Once cell locations were identified, single-cell fluorescence traces were extracted from corresponding locations in movies of unnormalized data blurred with 2d Gaussian filter with 1 pixel standard deviation.

#### Exclusion of spontaneously active and dying cells

While measuring a large number of cells in an acute slice, a portion of cells showed spontaneous activity, characterized by transient fluorescent increases uncorrelated with blue light stimulation; and cell death, characterized by a large and irreversible increase in fluorescence. For PCA analysis in **Fig. 3** and in **Fig. 4**, slices were imaged nine times, five times before AED application and four times after. Imaging epochs were averaged to generate movies with 108 frames (12 epochs per run x 9 runs). After extracting cell traces from these movies for all slices in the experiment, each cell’s mean and standard deviation per run were calculated. Least squares fits on the mean and standard deviation were performed on 3-pre drug runs and projected to the full nine runs. Cells were excluded from further analysis if any projected mean or standard deviation was less than 1/15 of the cell’s mean value or if the root mean square error of the fit was larger than 1/15 of the cell’s mean value. This procedure rejected < 17% of cells.

#### Generation of excitability maps

To generate the maps in **Fig. 3e** and **Supplementary Fig. 4j** the fluorescence trace for each included cell was normalized by subtracting its mean fluorescence values for each run and normalizing by the standard deviation for each run. For each cell, 3 pre-drug runs were averaged together to yield a 12 element vector corresponding to normalized F in each epoch. Principal component analysis yielded 3 main principal components which were then back-projected into pixel space for each slice, yielding the black and white images in **Supplementary Fig. 4j**. Color images were generated using L*a*b colorspace, by projecting PC1 into lightness, L, and PC2 and PC3 into the red-green and blue-yellow axes, *a* and *b*.

To generate maps of changes in drug response in **Fig. 4b**, ΔF images from four runs before and after drug addition were averaged together, median filtered with a 3 pixel kernel, saturated at their 99.5^th^ percentile, and displayed in the green and red channels, respectively. The blue channel is the average of the red and green images. Color saturation was adjusted in L*a*b space to aid in visualization. In **Fig. 5** ΔF images are scaled to the same absolute counts and shown in separate color channels.

#### Cortical layer analysis

All striatal cells were pooled and treated separately. For cortical cells, cortex boundaries were manually defined in structural images as the surface of the brain and the bottom of Layer 6. Boundaries were registered to functional images (above) and cells were assigned a normalized depth coordinate based on these boundaries. Drug response, defined as ΔF_drug_/ΔF_0_, could then be related to normalized cortical depth. For each slice, cells were binned by cortical depth and the drug response per cell averaged over cells. Extreme cell responses were excluded from each bin using the generalized extreme Studentized deviate test. Layer boundary locations were taken from the primary somatosensory cortex in the matched coronal slices of the Allen Brain Reference Atlas.

KCNQ3 expression levels were acquired from Allen Brain Institute experiment #100041071. The somatosensory cortex was manually defined in 11 sagittal slices from a P28 male mouse. The available expression image was used to mask the raw data, but expression values were obtained directly from the raw ISH data. The edges of the cortex and cortical depth bins were defined as above and expression values were averaged together across slices from the same experiment.

